# Population bottlenecks constrain host microbiome diversity and genetic variation impeding fitness

**DOI:** 10.1101/2021.07.04.450854

**Authors:** Michael Ørsted, Erika Yashiro, Ary A. Hoffmann, Torsten Nygaard Kristensen

## Abstract

It is becoming increasingly clear that microbial symbionts influence key aspects of their host’s fitness, and *vice versa*. This may fundamentally change our thinking about how microbes and hosts interact in influencing fitness and adaptation to changing environments. Here we explore how reductions in population size commonly experienced by threatened species influence microbiome diversity. Consequences of such reductions are normally interpreted in terms of a loss of genetic variation, increased inbreeding and associated inbreeding depression. However, fitness effects of population bottlenecks might also be mediated through microbiome diversity, such as through loss of functionally important microbes. Here we utilise 50 *Drosophila melanogaster* lines with different histories of population bottlenecks to explore these questions. The lines were phenotyped for egg-to-adult viability and their genomes sequenced to estimate genetic variation. The bacterial 16S rRNA gene was amplified in these lines to investigate microbial diversity. We found that 1) host population bottlenecks constrained microbiome richness and diversity, 2) core microbiomes of hosts with low genetic variation were constituted from subsets of microbiomes found in flies with higher genetic variation, 3) both microbiome diversity and host genetic variation contributed to host population fitness, 4) connectivity and robustness of bacterial networks was low in the inbred lines regardless of host genetic variation, 5) reduced microbial diversity was associated with weaker evolutionary responses of hosts in stressful environments, and 6) these effects were unrelated to *Wolbachia* density. These findings suggest that population bottlenecks reduce hologenomic variation (combined host and microbial genetic variation). Thus, while the current biodiversity crisis focuses on population sizes and genetic variation of eukaryotes, an additional focal point should be the microbial diversity carried by the eukaryotes, which in turn may influence host fitness and adaptability with consequences for the persistence of populations.

**Author summary:** It is becoming increasingly clear that organisms and the microbes that live on or in them – their microbiome – affect each other in profound ways that we are just beginning to understand. For instance, a diverse microbiome can help maintain metabolic functions or fight pathogens causing diseases. A disrupted microbiome may be especially critical for animals and plants that occur in low numbers because of threats from e.g. human exploitation or climate change, as they may already suffer from genetic challenges such as inbreeding and reduced evolutionary potential. The importance of such a reduction in population size, called a bottleneck, on the microbial diversity and the potential interactive effects on host health remains unexplored. Here we experimentally test these associations by investigating the microbiomes of 50 inbred or non-inbred populations of vinegar flies. We found that restricting the population size constrain the host’s genetic variation and simultaneously decreases the diversity of the microbiome that they harbor, and that both effects were detrimental to host fitness. The microbial communities in inbred host populations were less robust than in their non-inbred counterparts, suggesting that we should increasingly consider the microbiome diversity, which may ultimately influence the health and persistence of threatened species.

## Introduction

It is becoming increasingly clear that most eukaryotes live in intimate and complex relationships with microbial communities both in their external environment and on or within their body including their gut (McFall-Ngai *et al*. 2013). Numerous studies have documented how the presence and abundance of certain microbes can influence key aspects of host fitness, such as lifespan, fecundity, immune responses, metabolic health, behaviour, and thermal stress tolerance traits (Sommer & Bäckhed 2013; Ericsson & Franklin 2015; Mueller & Sachs 2015; Sampson & Mazmanian 2015; Sison-Mangus *et al*. 2015; Moghadam *et al*. 2018; Douglas 2019). Conversely, the host can also control microbial composition to some extent, such as through changing nutrient availability by diet choice or host metabolism (Muegge *et al*. 2011; Jehrke *et al*. 2018), triggering immune factors (Douglas 2019; Marra *et al*. 2021), or controlling the gut mechanically such as through peristalsis (Du *et al*. 2016). The genetic background of the host can also interact with the microbiome (Chandler *et al*. 2011; Dobson *et al*. 2015). For instance, host genes can affect the abundance of certain bacteria, allowing the microbial composition of hosts to be treated as a quantitative trait in genetic analysis (Chaston *et al*. 2016). These interactions between the host and its microbiota can in turn have a substantial impact on host fitness (Gould *et al*. 2018; Walters *et al*. 2019; West *et al*. 2019).

Because interactions between microbes and their hosts shape so many aspects of life, the impact on core biological processes including evolutionary adaptation may need to be reconsidered (Shapira 2016; Koskella & Bergelson 2020; Moeller & Sanders 2020; Henry *et al*. 2021). The ‘holobiont’ concept reflects the idea that eukaryote individuals do not act as autonomous units, but rather as networks consisting of the host and all its associated microbiota, and that their collective genomes – the ‘hologenome’ – forms a cohesive unit of selection (Zilber-Rosenberg & Rosenberg 2008; Bordenstein & Theis 2015; Kristensen *et al*. 2021). Some experimental support for this idea is emerging. For instance, host selection for thermal tolerance resulted in an altered microbial composition and modulated the microbes’ response to temperature (Kokou *et al*. 2018). While resident microbes respond to stressful environmental conditions, they can also subsequently aid the response of the host to such conditions (Moghadam *et al*. 2018). More generally, the microbial community may play a so-far underappreciated role in the broader context of population persistence which is important for research areas like conservation biology (Bahrndorff *et al*. 2016; Shapira 2016; Trevelline *et al*. 2019; West *et al*. 2019).

These conjectures raise the issue of how microbes respond to changes in a host’s population size. Populations that undergo repeated or persistent reductions in size frequently suffer from inbreeding depression and genetic drift resulting in lowered fitness and reduced genetic variations (Charlesworth & Charlesworth 1987; Kristensen & Sørensen 2005), ultimately impeding evolutionary capacity and increasing the risk of extinction (Spielman *et al*. 2004; Frankham 2005; Markert *et al*. 2010; Hoffmann *et al*. 2017; Ørsted *et al*. 2019; Willi *et al*. 2022). These patterns have been investigated from the perspective of the nuclear genetic background rather than the hologenome; for instance, inbreeding depression is normally assumed to reflect the expression of deleterious alleles in their homozygotic form. However, host population size decreases may also lead to microbial perturbations, with potential feed-back effects on host fitness which could contribute to additional lowering of population size (Redford *et al*. 2012; Bahrndorff *et al*. 2016; Trevelline *et al*. 2019). These interacting effects of microbial diversity, genetic diversity and fitness have not yet been widely tested.

Here we investigate microbial composition and abundance in 50 populations of *Drosophila melanogaster* that have experienced a variable number of generations of population bottlenecks. These populations were subsequently phenotyped for fitness components and sequenced to obtain a genome-wide molecular measure of genetic variation (Ørsted *et al*. 2019). This set of lines provides a unique resource to investigate associations between host genetic variation, host fitness and microbial diversity (**Fig. 1**). We consider two main questions. 1) How is microbiome richness and diversity associated with host genetic variation imposed by population bottlenecks? 2) Is both host microbiome diversity and genetic variation associated with host fitness, and if so, do they contribute independently? Answers to these questions increase our understanding of the importance of microbiomes in conservation genetics and evolutionary biology. We hypothesise that reduced host genetic variation provides a less diverse environment for microbes leading to reduced microbial diversity through fewer host genotype-microbial taxon-specific interactions (**Fig. 1**). Evidence suggests that increased microbiome diversity is adaptive for the host and that increased hologenome genetic variation facilitates faster evolutionary responses to selection (Hauffe & Barelli 2019; Henry *et al*. 2021; Kristensen *et al*. 2021). Our work further highlights the importance of considering both host and microbiome diversity for predicting evolutionary consequences of population bottlenecks in small and threatened populations.

**Figure 1.**
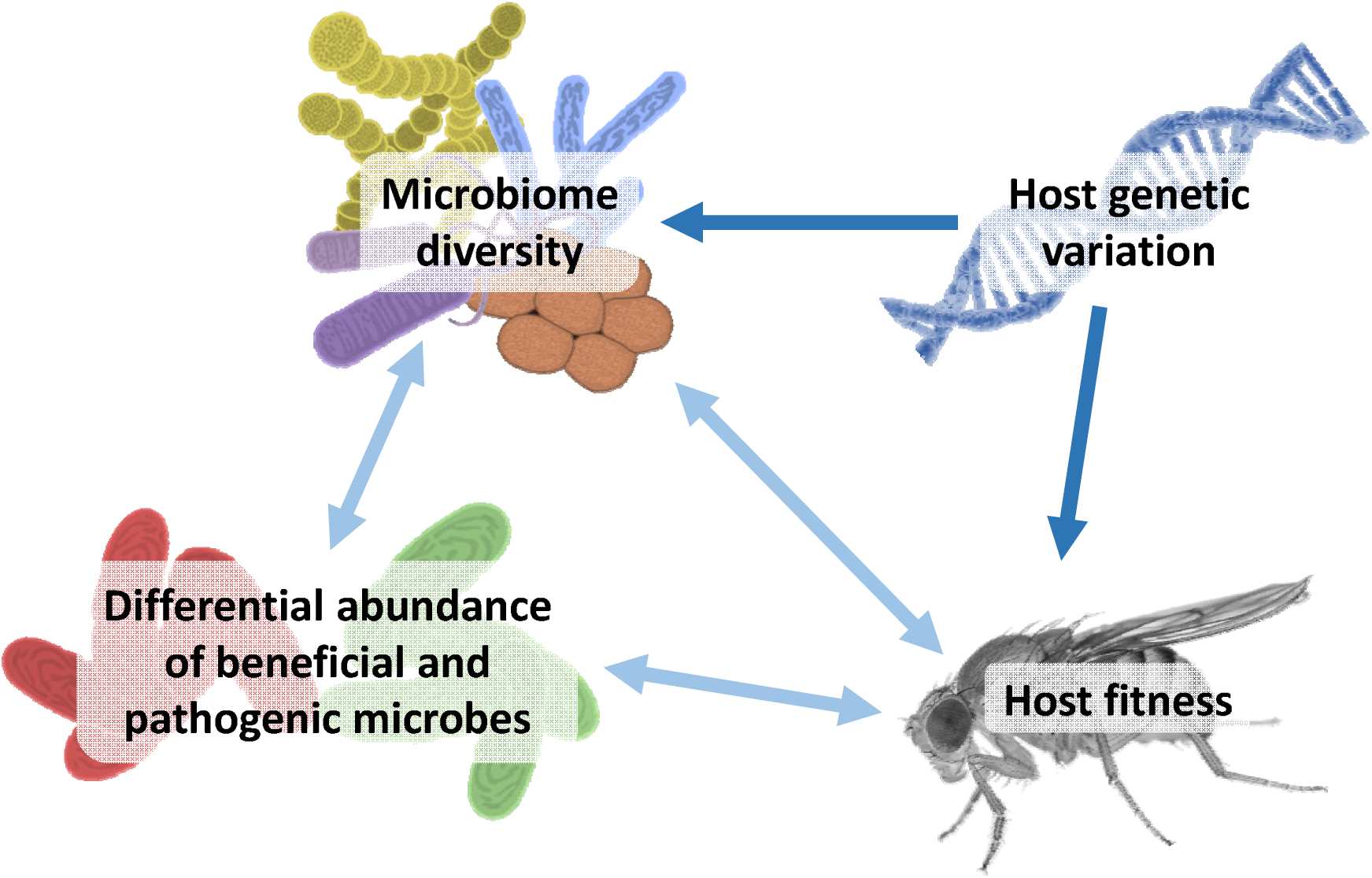
Hypothetical associations between host genetic variation, microbiome diversity and host fitness. Conceptual illustration of hypothetical associations between host genetic variation, microbiome diversity and host fitness in the experimental set of lines. Host genomic variation was manipulated through exposing populations to a variable number of bottlenecks which is assumed to have a causal effect on fitness and microbial diversity. The dark blue arrows represent potential unidirectional effects, and light blue arrows represent potential bidirectional effects. In one line of causality, low genetic variation in the host constrains the diversity (species richness and abundance) of the host-associated microbiome, which in turn affects host fitness directly, i.e. through supporting fewer host genotype-microbial taxon-specific interactions that are functionally important, and/or indirectly through changing the composition and relative abundance of beneficial or pathogenic microbes. In another line of causality, low host genetic variation directly affects host fitness which in turn leads to a less diverse microbiome through physiological or metabolic changes in the host “environment”, or indirectly such as through a higher abundance of detrimental pathogenic microbes resulting from a weakened immune response.

## Results

We selected 50 lines for microbiome analysis, which represent a subset of 109 inbred lines that have been through marked reductions in population size for a varying number of generations as well as 10 outbred control lines kept at a large population size (for details see Ørsted et al. 2019). These original total 119 lines had been sequenced using Genotyping-By-Sequencing (GBS) to allow nucleotide diversity (π) to be estimated. In addition, the fitness component egg-to-adult viability (hereafter just viability) was determined in each line (see Ørsted *et al*. 2019 for details on genomic variation and phenotypes assessed). We found a positive correlation between viability and π for these 119 lines (r_s_=0.45; *p* < 0.001; **Supplementary Fig. S1**) indicating inbreeding depression for viability which is used here as a proxy for Darwinian fitness. For the microbiome characterization, two groups each consisting of 25 lines were selected based on measures of standardized viability and π (sum of Z-scores, see methods). One group of 25 lines had low genetic variation and low viability, and another group of 25 lines had high genetic variation and high viability. The ‘high’ group included nine of the outbred controls as the outbred lines generally had very high viability and nucleotide variability (**Supplementary Table S1**). The inbreeding procedure resulted in varying degrees of nucleotide diversity in the resulting lines, and as such the genetic variation of the outbred lines and the inbred lines in the ‘high’ group did not significantly differ (**Supplementary Fig. S2A**). However, to distinguish between effects of genetic variation within inbred lines from the effects of bottlenecks itself (inbreeding/outbreeding), we separated lines into three categories: ‘low genetic variation’ and ‘high genetic variation’ inbred lines and ‘outbred’ (OB) control lines (**Supplementary Fig. S2**). Six lines evenly distributed among the genetic variation groups were removed due to a high relative abundance of the microbial endosymbiont *Wolbachia*, which might affect host fitness and/or abundance of other microbiome taxa, resulting in 44 being analysed (see methods for details).

### Loss of host genomic variation decreases microbiome richness and diversity

Loss of host genetic variation was associated with decreased microbiome richness and diversity. The microbiomes from the outbred flies and the high genetic variation lines had a higher level of community diversity than the microbiomes of flies from the low genetic variation group (**Fig. 2**). Generally, we observed an increase in microbiome diversity with increasing fly host genetic variation and viability, regardless of which measure we used to quantify the microbiome diversity. Richness indices, namely observed amplified sequence variant (ASV) richness and chao1 estimated ASV richness (**Fig. 2A-B**), showed a stronger trend than the diversity indices accounting for relative abundances (Shannon-Wiener and Simpson’s; **Fig. 2C-D**). Even at the class level, the microbiomes from the low genetic variation group were less taxonomically heterogeneous than those from the high genetic variation and outbred groups (**Supplementary Fig. S3**). Most bacteria in the *D. melanogaster* microbiomes belonged to the Alphaprotoebacteria, Bacilli, and to a lesser extent Bacteroidia and Actinobacteria. The increasing level of taxonomic diversity with increasing host genetic variation was further evident at the genus level, with the low genetic variation group harbouring the fewest number of genera, while the outbred flies had the highest number (**Supplementary Fig. S4**). Non-metric multidimensional scaling (NMDS) analysis based on the weighted and unweighted UniFrac distances (**Fig. 3**) showed that the microbiomes differed depending on the fly host’s level of genetic variation. Generally, there was better segregation of the fly groups when only the microbiome membership (unweighted UniFrac) was considered (**Fig. 3C-D)**, with a clear trend across the viability and nucleotide diversity gradient across the first dimension of ordination space. Meanwhile the relative abundance patterns of the ASVs across the lines created noise, notably among the high and outbred groups (**Fig. 3A-B)**. However, the community membership was more similar among the outbred lines and more dispersed within the inbred lines (**Fig. 3C-D**), while the abundance patterns showed greater similarity among the low genetic variation flies than the higher variation and outbred flies, due to the high relative abundance of a few ASVs in common in the low variation flies (**Fig. 3A-B**). PERMANOVA analysis showed that the fly groupings explained 22.8% and 43.0% of the microbial community membership and structure (i.e. membership + relative abundance using weighted UniFrac), respectively (*p* < 0.001). The host traits, nucleotide diversity and viability were significantly associated with the microbiome community patterns (significant envfit arrows, **Fig. 3A-D**). Viability explained a greater proportion of the microbiome composition than did nucleotide diversity, where nucleotide diversity and viability explained 15.9% and 17.8% of the community membership, while explaining 27.7% and 35.3% of the community structure.

**Figure 2.**
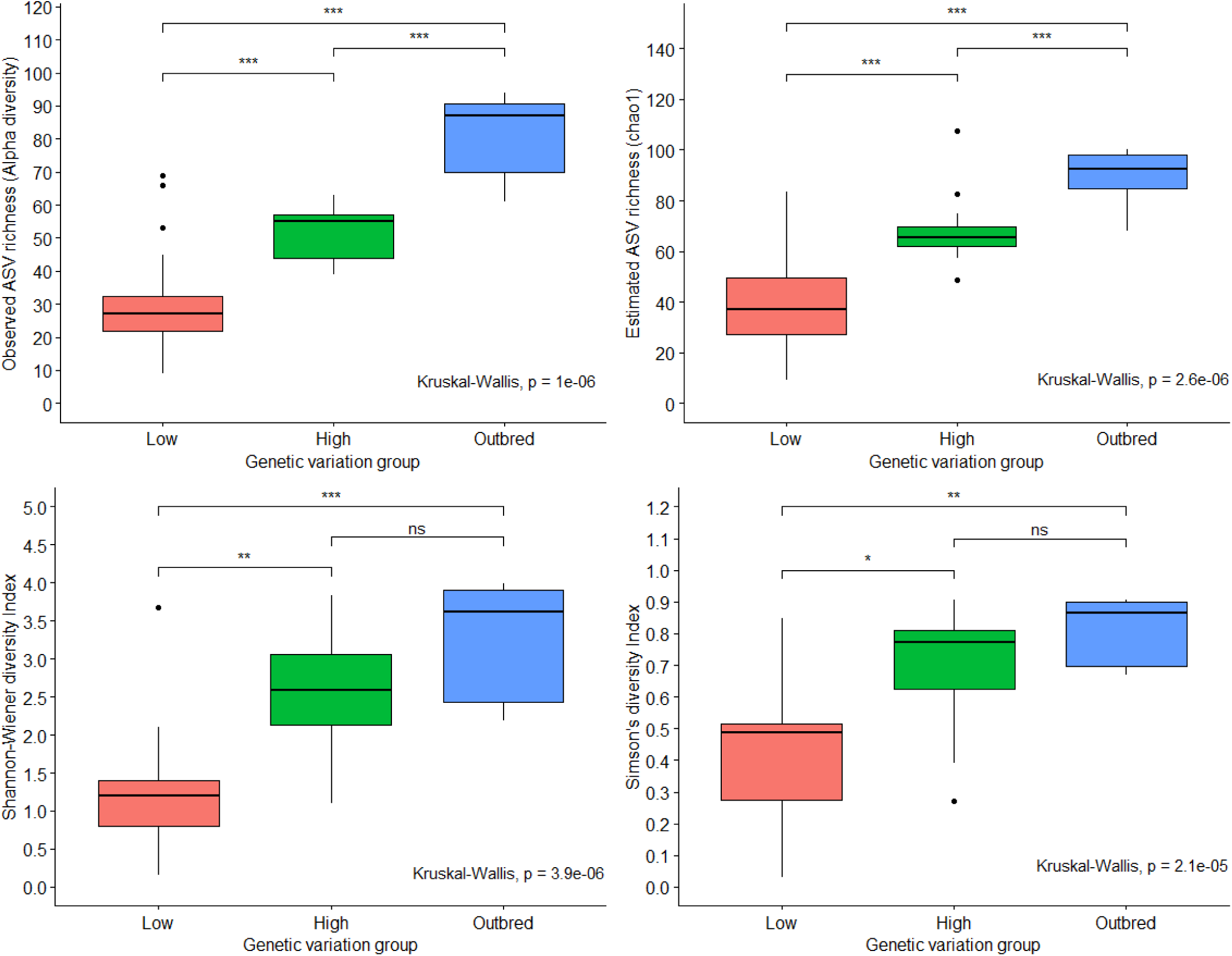
Host population bottlenecks decrease microbiome diversity. Boxplots of microbiome diversity in the three host genetic variation groups; low genetic variation (Low; red), high genetic variation (High; green), and outbred (blue) measured as either richness indices (**A**. observed ASV richness (Alpha richness), and **B**. estimated ASV richness (chao1 estimation)) or measured using indices accounting for individual ASV abundances as well (**C**. Shannon-Wiener Index, and **D**. Simpson’s Index). In all panels, the *p* values of a Kruskal-Wallis test show a significant effect of group, while asterisks denote the results of pairwise Wilcox’s t-tests between groups; *** *p* < 0.001; ** *p* < 0.01; * *p* < 0.05; and ns: *p* > 0.05.

**Figure 3.**
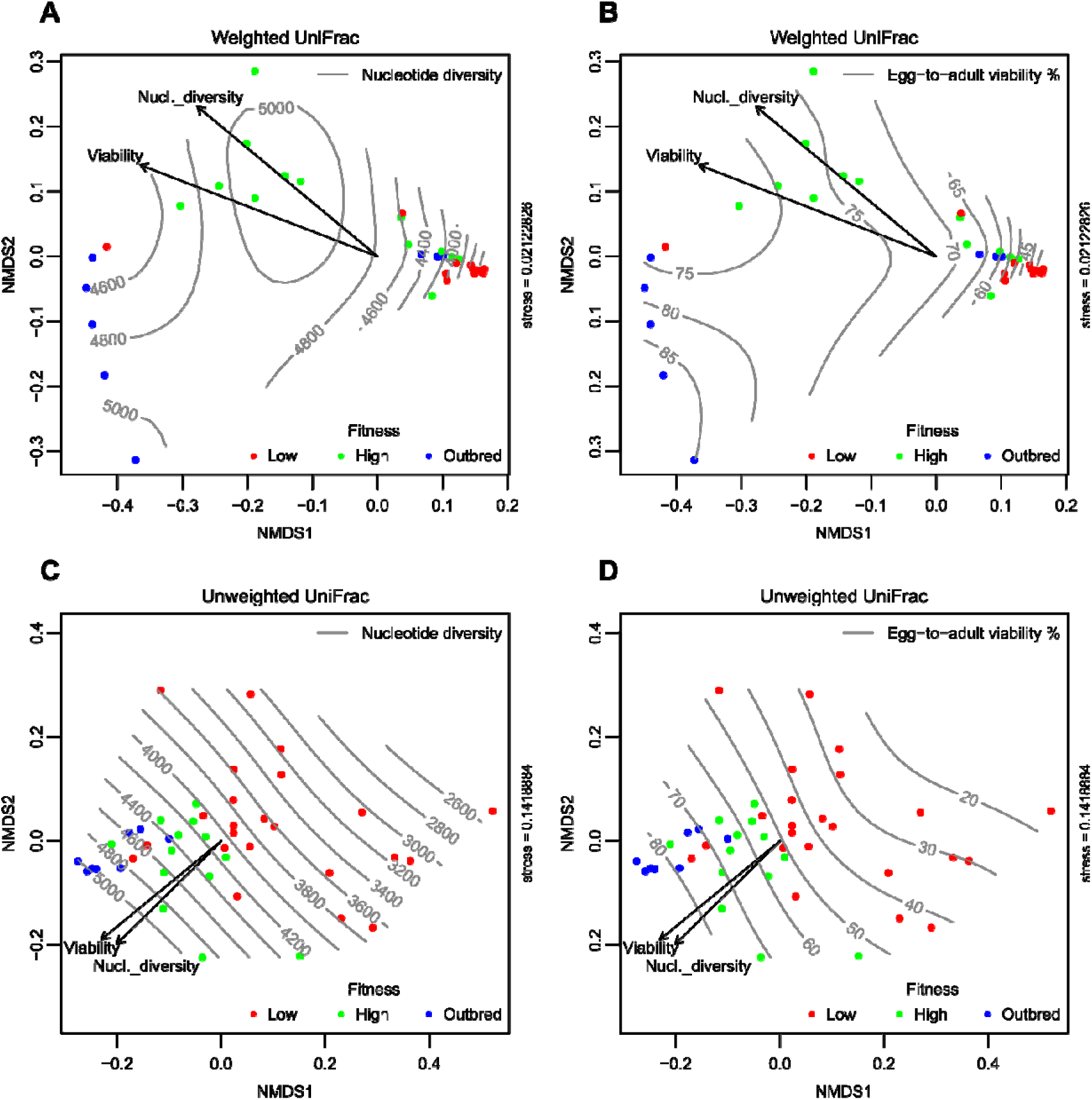
Microbiome composition differ depending on the host’s level of genetic variation. Non-metric multidimensional scaling (NMDS) plots based on the UniFrac distances (weighted; **A-B**, and unweighted; **C-D**) between the 44 *D. melanogaster* lines (low genetic variation (red), high genetic variation (green), and outbred (blue). UniFrac distances account for the relative relatedness of community members, where weighted unifrac incorporates the abundance of observed organisms, while unweighted unifrac only considers presence or absence. Isolines of associated covariates are shown for nucleotide diversity (**A** and **C**) and egg-to-adult viability (**B** and **D**). Envfit values of significant drivers (*p* < 0.05) are shown as arrows (viability and nucleotide diversity, respectively).

### Both host genetic variation and microbiome diversity contribute to host fitness

To identify drivers of host fitness, we fitted generalized linear mixed effects models (GLMMs) with egg-to-adult viability as a function of host nucleotide diversity (π) and microbiome diversity and their interaction (one model for each of the four different measures of microbiome diversity; see methods). For all microbiome diversity metrics, both π and microbiome diversity contributed to host fitness, with π contributing to a greater extent (∼2x larger scaled effect sizes; **Table 1; Supplementary Table S2**). For Alpha richness, we observed a significant positive interaction between host genetic variation and microbial diversity on host fitness, meaning that they did not act independently. In fact, the positive interaction coefficients suggested synergism; i.e. in lines with high genetic variation, the effect of increasing microbiome richness was greater than in lines with low genetic variation and *vice versa* (**Supplementary Fig. S5**), while for chao1 ASV richness, Shannon-Wiener and Simpson’s indices, host genetic variation and microbial diversity contributed independently to host fitness (**Supplementary Table S2)**. In all cases, the full models had higher explanatory power than each individual model (π or microbiome diversity alone), based on χ^2^ tests (**Table 1; Supplementary Table S2**). Using data from Ørsted *et al*. (2019) for the set of lines used in the present study, we also associated microbial diversity with evolutionary responses in two traits (dry body mass and productivity measured as eggs per female per day) here defined as the slope of an ordinary linear regression across 10 generations of rearing on a stressful medium. Interestingly, we found an effect of microbial diversity on evolutionary responses for both traits (**Supplementary Fig. S6**). For details on assessment of these phenotypes, see Ørsted *et al*. (2019).

**Table 1.**
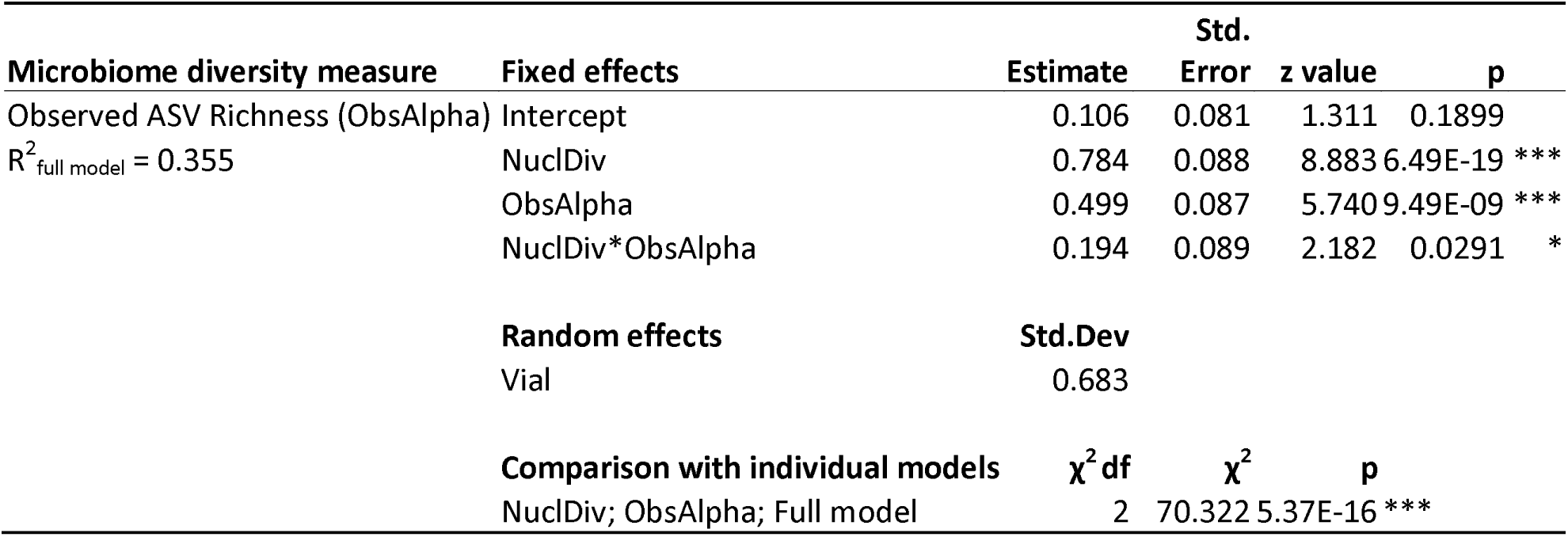
Host genetic variation and microbiome richness interact synergistically on host fitness.

Results of the general linear mixed model (GLMMs) of egg-to-adult viability as a function of nucleotide diversity (NuclDiv) and microbiome diversity (Alpha richness; ObsAlpha) and their interaction as fixed effects. Both dependent and independent variables are scaled (Z-standardization) to allow direct comparison of effect sizes. Replicate vial IDs were included as a random effect, as flies from the same vial are not considered independent. Conditional coefficients of determination of the GLMMs 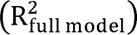 interpreted as the variance explained by the entire model, including both fixed and random effects, are shown. Asterisks denote the significance of individual variables or interactions; *** *p* < 0.001; ** *p* < 0.01; and * *p* < 0.05. The full model including both dependent variables and their interaction is compared with individual models with either nucleotide diversity or alpha diversity with a χ^2^ test.

### Host genetic variation is associated with differential bacterial relative abundance

The subset of the bacterial taxa that were differentially more abundant in the outbred flies were more diverse than in the inbred lines (i.e. the low and high genetic variation groups), with as many as 54 ASVs belonging to five bacterial classes and 12 orders (**Supplementary Table S3**). In comparison, only one *Acetobacter* ASV was significantly more abundant in the bottlenecked flies, i.e. the low and high genetic variation groups. More generally, at both the ASV and genus levels, *Acetobacter* relative abundances were particularly high in the low genetic variation lines compared to the lines with higher genetic variation (**Fig. 4A-C**), with *Acetobacter* ASVs making up the majority of the communities in the low genetic variation lines (median 94.0% for ASVs 1+2 and 96.9% for the genus). In contrast, five ASVs belonging to *Lactobacillus* and *Enterococcus* were significantly more abundant in the high genetic variation and outbred flies, while at the genus level, *Enterococcus* but not *Lactobacillus* showed this same trend (**Fig. 4 D-F**, and **Supplementary Table S3**). In addition, certain ASVs displayed incremental increases in relative abundances with increasing level of genetic variation and outbreeding, despite DESeq results being non-significant (**Fig. 4F-H**). Since relative abundances are used throughout the present study, these differential abundances do not correspond to simple changes in diversity, but actual changes in certain ASVs shifting between genetic variation groups.

**Figure 4.**
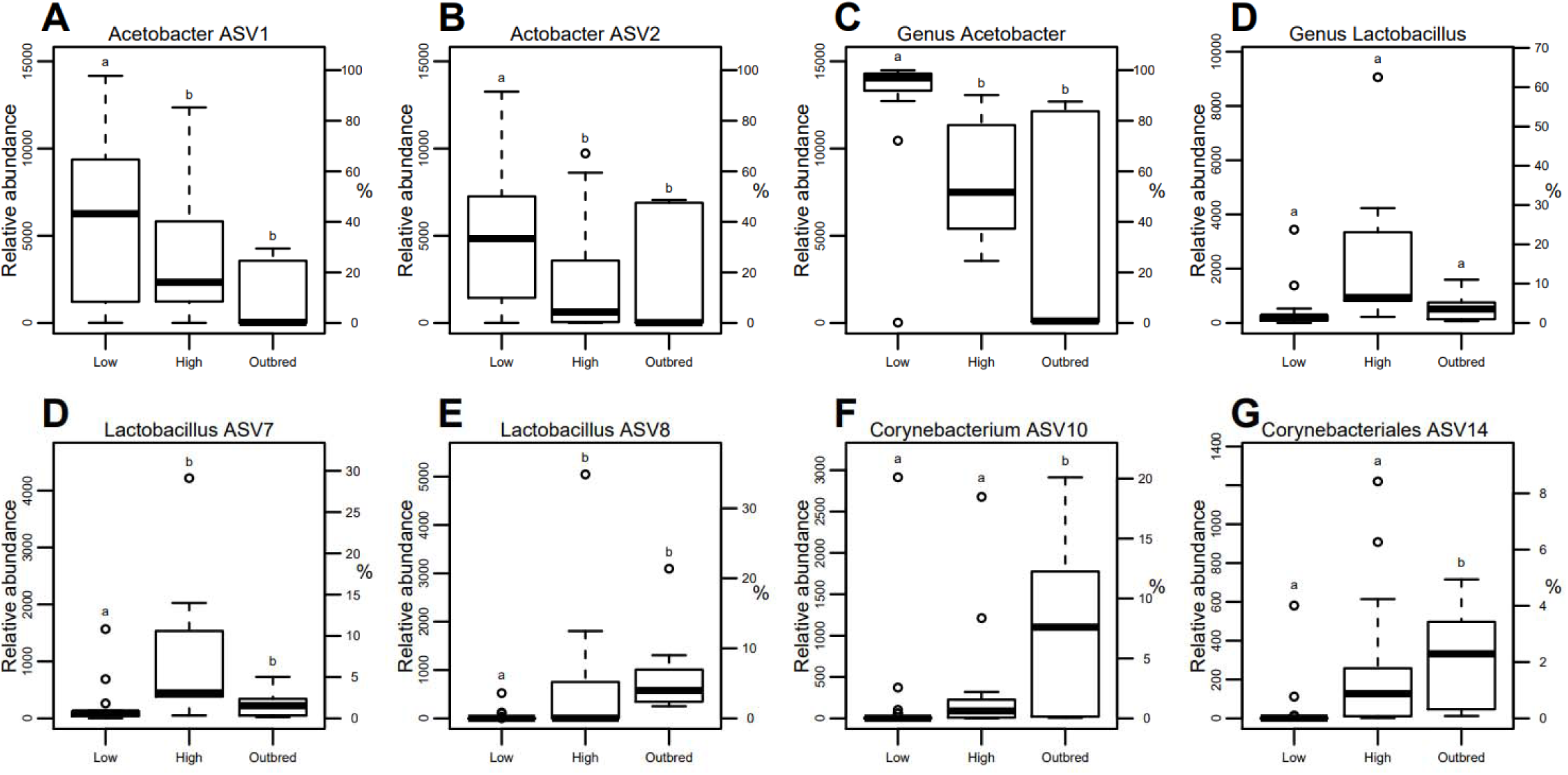
Host genetic variation is associated with differential abundance of microbial taxa. Relative abundance of differentially enriched bacterial ASVs and genera **A-C.** *Acetobacter* ASVs and genus, **D-E.** *Lactobacillus* ASVs and genus, and **F-G.** Corynebacteriales ASVs in the three host genetic variations groups; low genetic variation (Low), high genetic variation (High), and outbred. The abundance axes are not scaled the same for all of the ASVs and genera because *Acetobacter* constitute the majority of the microbiome of the low genetic variation group. Letters denote significant differences in relative abundance of ASVs or genera between fly groups (DESeq2 Wald test adjusted *p* < 0.05). The percent reads are relative to the number to which all lines were initially rarefied (14,488 reads/sample).

### Core microbiomes of the low genetic variation lines are subsets of the high genetic variation lines

The number of ASVs that were present in at least 80% of lines was defined as the core microbiome. The number of ASVs belonging to this core decreased with decreasing host genetic diversity in the fly groups, with 48, 22, and 11 ASVs in outbred, high, and low genetic variation flies respectively (**Table 2**). Eleven of these ASVs persisted as part of the core microbiome across all of the fly groups, and belonged to the *Acetobacter*, *Enterococcus*, *Lactobacillus*, *Leuconostoc*, and Bacilli. Meanwhile, nine ASVs belonging to the *Corynebacterium*, *Empedobacter*, Nocardiaceae*, Enterococcus*, *Mesorhizobium*, and *Lactobacillus*, were only found in the core microbiomes of the high and outbred flies. The core microbiome of the lower genetic variation lines were subsets of those present in the higher genetic variation lines and outbreds, with 11/11 low genetic variation line ASVs represented in the high genetic variation lines, and 20/22 high genetic variation line ASVs represented in the outbred lines (**Fig. 5, Table 2**).

**Figure 5.**
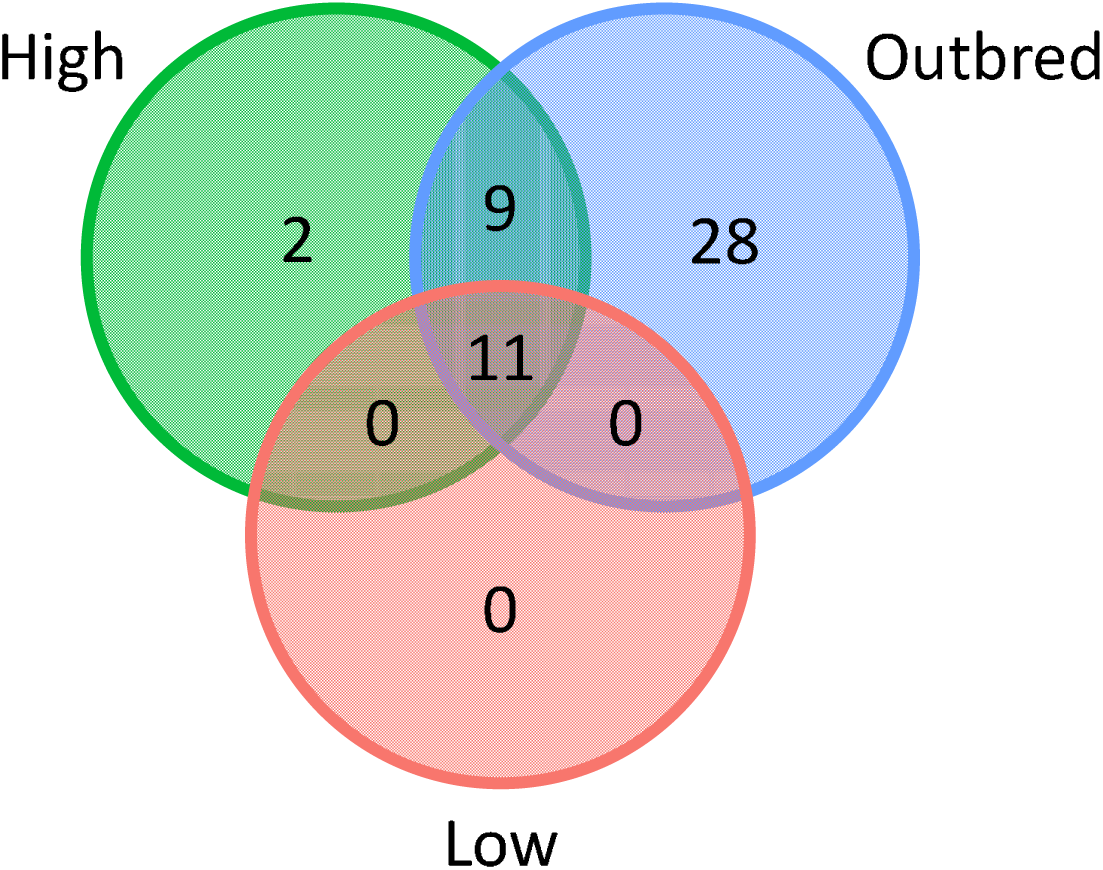
Core microbiomes of the low genetic variation lines are subsets of the high genetic variation lines. Venn-diagram showing that the core microbiomes of the low genetic variation lines (Low; red) are subsets of those present in the higher genetic variation lines (High; green) and outbred lines (blue), with 11/11 low genetic variation group ASVs represented in the high genetic variation group, and 20/22 high genetic variation group ASVs represented in the outbred group. The taxonomy of individual ASVs can be seen in **Table 2**.

**Table 2.** Core microbiomes of the three host genetic variations groups. Core microbiome ASVs in each of the three host genetic variations groups: low genetic variation (Low), high genetic variation (High), and outbred (OB). The ASVs belonging to the core microbiome for each fly group is represented by + (core) and – (not core). Here we define presence in the core microbiome if an ASV is present in at least 80% of lines of a particular group. The lowest taxonomy level is listed (o__: order, f__: family, g__:genus).

| ASVs | Class | Order | Family | Genus | Species | Low | High | OB |
| --- | --- | --- | --- | --- | --- | --- | --- | --- |
| ASV1 | Alphaproteobacteria | Acetobacterales | Acetobacteraceae | *Acetobacter* | *sp.* | + | + | + |
| ASV2 | Alphaproteobacteria | Acetobacterales | Acetobacteraceae | *Acetobacter* | *sp.* | + | + | + |
| ASV4 | Alphaproteobacteria | Acetobacterales | Acetobacteraceae | *Acetobacter* | *sp.* | + | + | + |
| ASV5 | Bacilli | Lactobacillales | Enterococcaceae | *Enterococcus* | *sp.* | + | + | + |
| ASV7 | Bacilli | Lactobacillales | Lactobacillaceae | *Lactobacillus* | *plantarum* | + | + | + |
| ASV8 | Bacilli | Lactobacillales | Enterococcaceae | *Enterococcus* | *sp.* | - | - | + |
| ASV9 | Bacilli | Lactobacillales | Leuconostocaceae | *Leuconostoc* | *sp.* | + | + | + |
| ASV10 | Actinobacteria | Corynebacteriales | Corynebacteriaceae | *Corynebacterium* | *sp.* | - | + | + |
| ASV11 | Bacteroidia | Flavobacteriales | Weeksellaceae | *Empedobacter* | *sp.* | - | + | + |
| ASV12 | Bacilli | Lactobacillales | Lactobacillaceae | *Lactobacillus* | *sp.* | + | + | + |
| ASV13 | Actinobacteria | Corynebacteriales | Nocardiaceae | *g__* | *sp.* | - | + | + |
| ASV14 | Actinobacteria | Corynebacteriales | *f__* | *g__* | *sp.* | - | + | + |
| ASV15 | Bacilli | Lactobacillales | Lactobacillaceae | *Lactobacillus* | *sp.* | + | + | + |
| ASV16 | Bacilli | Lactobacillales | Lactobacillaceae | *Lactobacillus* | *plantarum* | + | + | + |
| ASV17 | Actinobacteria | Corynebacteriales | Nocardiaceae | *g__* | *sp.* | - | + | + |
| ASV18 | Bacilli | Lactobacillales | Enterococcaceae | *Enterococcus* | *sp.* | - | + | + |
| ASV19 | Actinobacteria | Corynebacteriales | Corynebacteriaceae | *Corynebacterium* | *sp.* | - | - | + |
| ASV21 | Alphaproteobacteria | Caulobacterales | Caulobacteraceae | *Brevundimonas* | *sp.* | - | - | + |
| ASV22 | Bacteroidia | Sphingobacteriales | Sphingobacteriaceae | *Sphingobacterium* | *mizutaii* | - | - | + |
| ASV23 | Gammaproteobacteria | Burkholderiales | Alcaligenaceae | *Alcaligenes* | *sp.* | - | - | + |
| ASV24 | Bacteroidia | Sphingobacteriales | Sphingobacteriaceae | *Sphingobacterium* | *mizutaii* | - | - | + |
| ASV25 | Alphaproteobacteria | Rhizobiales | Rhizobiaceae | *Mesorhizobium* | *sp.* | - | + | + |
| ASV27 | Bacilli | Lactobacillales | Lactobacillaceae | *Lactobacillus* | *brevis* | + | + | + |
| ASV28 | Gammaproteobacteria | Burkholderiales | Alcaligenaceae | *Alcaligenes* | *sp.* | - | - | + |
| ASV29 | Actinobacteria | Corynebacteriales | Nocardiaceae | *Rhodococcus* | *sp.* | - | - | + |
| ASV30 | Actinobacteria | Micrococcales | Microbacteriaceae | *Leucobacter* | *sp.* | - | - | + |
| ASV32 | Alphaproteobacteria | Rhizobiales | Rhizobiaceae | *g__* | *sp.* | - | - | + |
| ASV36 | Actinobacteria | Micrococcales | Micrococcaceae | *Glutamicibacter* | *sp.* | - | - | + |
| ASV40 | Actinobacteria | Micrococcales | Microbacteriaceae | *Microbacterium* | *sp.* | - | - | + |
| ASV45 | Alphaproteobacteria | Caulobacterales | Caulobacteraceae | *Brevundimonas* | *sp.* | - | - | + |
| ASV46 | Actinobacteria | Micrococcales | Brevibacteriaceae | *Brevibacterium* | *Brevibacterium* | - | - | + |
| ASV48 | Bacilli | Bacillales | Planococcaceae | *Lysinibacillus* | *sp.* | - | - | + |
| ASV53 | Bacilli | Lactobacillales | Enterococcaceae | *Enterococcus* | *sp.* | - | + | + |
| ASV56 | Bacilli | Lactobacillales | Lactobacillaceae | *Lactobacillus* | *brevis* | - | + | + |
| ASV68 | Bacilli | Lactobacillales | Enterococcaceae | *Enterococcus* | *sp.* | - | - | + |
| ASV72 | Alphaproteobacteria | Rhizobiales | Rhizobiaceae | *Mesorhizobium* | *sp.* | - | - | + |
| ASV89 | Bacilli | Lactobacillales | Enterococcaceae | *Enterococcus* | *sp.* | - | - | + |
| ASV91 | Actinobacteria | Corynebacteriales | Corynebacteriaceae | *Corynebacterium* | *flavescens* | - | - | + |
| ASV93 | Bacilli | Bacillales | Planococcaceae | *Lysinibacillus* | *sp.* | - | - | + |
| ASV94 | Bacilli | Lactobacillales | Lactobacillaceae | *Lactobacillus* | *sp.* | - | + | - |
| ASV105 | Bacilli | Bacillales | Planococcaceae | *Lysinibacillus* | *sp.* | - | - | + |
| ASV108 | Bacilli | Bacillales | Planococcaceae | *Lysinibacillus* | *sp.* | - | - | + |
| ASV110 | Bacilli | Bacillales | Planococcaceae | *Lysinibacillus* | *sp.* | - | - | + |
| ASV114 | Bacilli | Bacillales | Planococcaceae | *Lysinibacillus* | *sp.* | - | - | + |
| ASV115 | Bacilli | Bacillales | Planococcaceae | *Lysinibacillus* | *sp.* | - | - | + |
| ASV117 | Bacilli | Bacillales | Planococcaceae | *Lysinibacillus* | *sp.* | - | - | + |
| ASV152 | Bacilli | *o__* | *f__* | *g__* | *sp.* | + | + | + |
| ASV189 | Bacilli | Lactobacillales | Enterococcaceae | *Enterococcus* | *sp.* | - | - | + |
| ASV193 | *c__* | *o__* | *f__* | *g__* | *sp.* | - | - | + |
| ASV383 | Bacilli | Lactobacillales | Leuconostocaceae | *Leuconostoc* | *sp.* | - | + | - |

### Bacterial co-abundance networks decreased in complexity with lower host genetic variation

In parallel to the DESeq analysis, which highlighted ASVs and genera, which were categorically enriched in the different fly lines, co-abundance network analysis across the microbiome dataset was performed to identify bacterial groups that were co-varying in relative abundance with the host fitness traits of the lines and co-varying with each other (**Fig. 6**). Fly egg-to-adult viability and nucleotide diversity co-varied with a large cluster of bacterial ASVs within the same modular cluster (**Fig. 6A**). Generally, viability co-varied with a larger group of bacterial ASVs than nucleotide diversity, giving 40 and 13 associations, respectively, while 12 of these ASVs co-varied with both viability and nucleotide diversity. Interestingly, most of the ASVs that co-varied with viability were either differentially enriched in the outbred flies or in both the outbred and high genetic variation flies (**Fig. 6B**). The ASVs that were differentially enriched in both outbred and high genetic variation flies belonged to 24 different taxonomic groups (**Fig. 6B, Supplementary Fig. S7A**). In contrast, the two *Acetobacter* ASVs that were significantly enriched in the low genetic variation flies belonged to separate modular clusters as highly prevalent negatively correlated ASVs (**Supplementary Fig. S7A**). The smaller modules were often exclusively made up of related ASVs, notably modules with *Lactobacillus* and *Enterococcus* species. The networks of each individual fly group (**Fig. 6C,D,E**) revealed that inbreeding the flies resulted in a lower number of co-varying ASVs, and decreased the degree of connectivity (i.e. the number of significant correlations) and overall complexity of the network, with the latter consisting of only a few ASVs in a modular structure. Having a higher host genetic variation did not result in a higher degree of covariation among ASVs among inbred lines. Host genetic variation was both positively and negatively correlated with the abundance of bacteria in the outbred network, while host genetic variation did not significantly co-vary with bacterial abundance in the high and low genetic variation networks (at correlation adjusted *p* < 0.05). Moreover, the ASVs belonging to the *Acetobacter*, that were significantly enriched in the low genetic variation flies, displayed covariation only with *Lactobacillus* among the inbred flies, (Fig. 6E), in contrast to those ASVs in the outbred flies.

**Figure 6.**
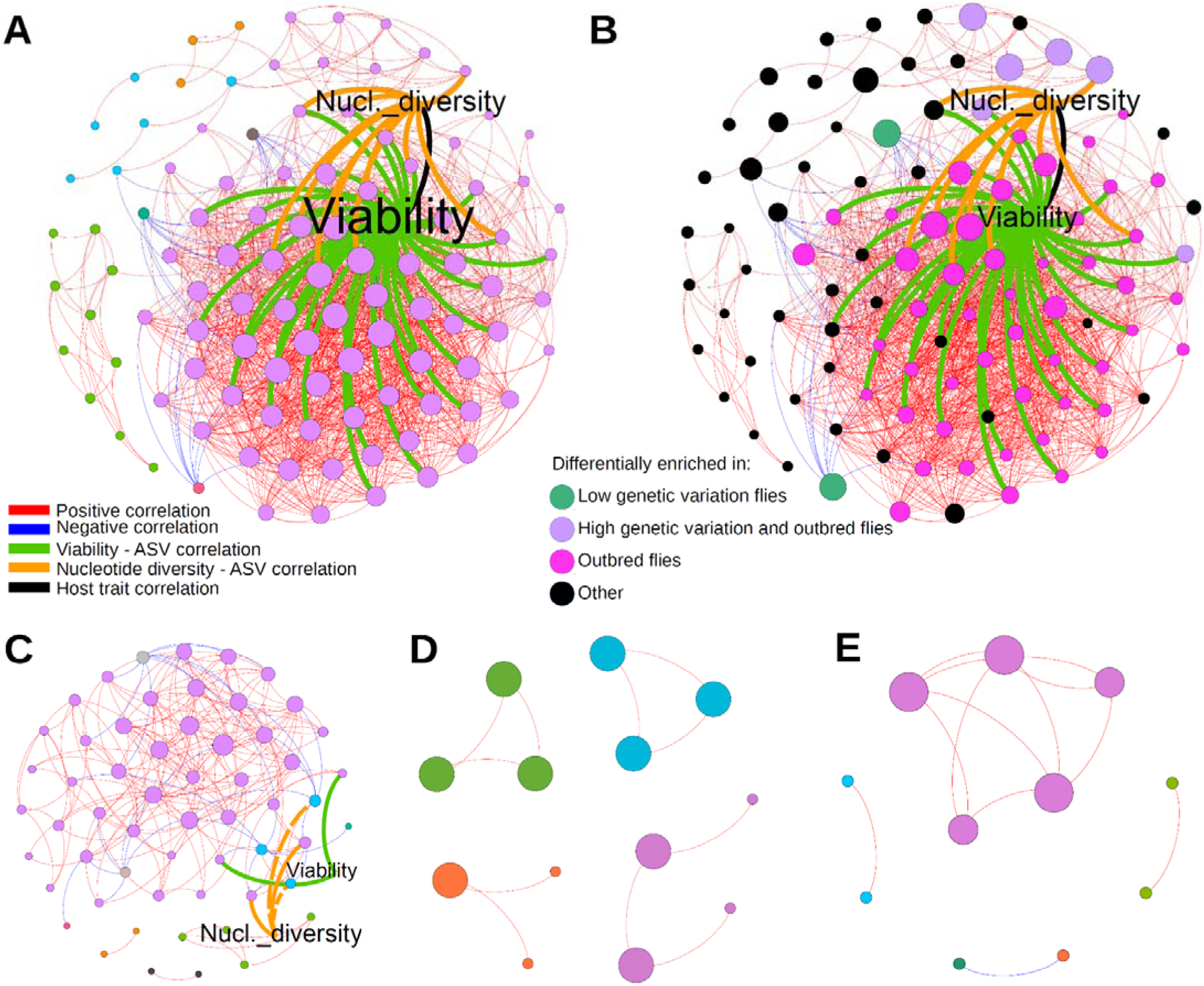
Bacterial co-abundance networks decreases in complexity with lower host genetic variation. Co-abundance networks of the fly microbiome. ASVs present in at least 30 reads in total and in at least three lines; correlations > 0.5 (or <-0.5) and fdr-corrected p-values < 0.05 are shown. The nodes are individual ASVs and the host fitness traits, egg-to-adult viability (Viability) and nucleotide diversity (Nucl. diversity), while the edges represent positive and negative correlations, and correlations linking host fitness traits and bacterial ASVs (which were positive correlations). The network containing lines from all of the fly groups (**A, B**), the outbred (**C**), high genetic variation (**D**), and low genetic variation (**E**) groups are shown. In **A, C, D, E,** node sizes display the degree of connectivity (relative within each figure) and colors mark the modularity structure (i.e. in which community, or cluster, the ASVs belong to based on the Leiden algorithm). The orange and green edges connecting Viability and Nucleotide diversity were positive except for two instances in **C** where the negative correlations are displayed with dotted lines. In **B**, node sizes display the number of lines within which the ASVs are present, and colors mark the major groups in which the ASVs are differentially enriched. The taxonomic assignment of each node can be seen in **Supplementary Fig. S7**.

## Discussion

In this study, we investigated the effects of host genetic variation on the microbiome diversity in *D. melanogaster* lines with experimentally manipulated levels of genetic variation. Consistent with a growing body of literature (Sommer & Bäckhed 2013; Ericsson & Franklin 2015; Mueller & Sachs 2015; Sampson & Mazmanian 2015; Sison-Mangus *et al*. 2015; Moghadam *et al*. 2018; Douglas 2019), we show that the microbial community of the host is strongly linked to the fitness of the host, in our case the viability of offspring. In addition, we link reductions in host genetic variation with microbiome diversity and clearly show that microbiome diversity is lower in lines with reduced genetic variation, regardless of whether we use richness or relative abundance metrics (**Fig. 2**). Only a few previous studies have investigated the association between components of the microbiota and host genetics in invertebrates, and those that did have mainly focused on genetic variability in the host’s ability to control the presence and/or abundance of specific endosymbionts (e.g., Mouton *et al*. 2007; von Burg *et al*. 2008; Chong & Moran 2016; Simhadri *et al*. 2017). Conversely, a few studies have investigated how endosymbionts can affect genetic variation in the host by distorting sex-ratios, such as through endosymbiont induction of cytoplasmic incompatibility or parthenogenesis (Engelstädter & Hurst 2009), potentially decreasing the effective population size and increasing genetic drift (Feldhaar 2011; Ferrari & Vavre 2011) or by influencing population dynamics and dispersal possibly affecting host genetic variation (Leonardo & Mondor 2006; Goodacre *et al*. 2009).

Despite recent calls for an integration of microbiome research in evolutionary and conservation biology (Bahrndorff *et al*. 2016; Grueber *et al*. 2019; Hauffe & Barelli 2019; Trevelline *et al*. 2019; West *et al*. 2019; Cullen *et al*. 2020; Moeller & Sanders 2020; Henry *et al*. 2021), little progress has been made on experimentally testing fundamental associations between population size, host genetic variation and microbial diversity. Here we provide a novel demonstration that restricting host population size has a profound impact on both the richness and the diversity of host-associated microbiomes, and that effects of host genetic variation and microbial richness interact synergistically in affecting host fitness. We also provide evidence that increased microbial diversity in the lines is associated with a stronger evolutionary response to stressful environments compared to responses seen in lines with less microbial diversity (**Supplementary Fig. S6**). Thus, we show that populations with high host genomic variation harbor the most diverse microbial community, and *vice versa* for populations with low genetic variation, and that these lines are least resilient to evolutionary pressure. According to the hologenome hypothesis, the holobiont constitutes a single unit of selection, thus our results suggest that small and fragmented populations have a reduced potential for responding to selection pressures not only due to reduced genetic variation and high rates of inbreeding but also due to reduced microbiome variation, effectively reducing hologenomic variation. Maintaining a diverse microbiome community may therefore be crucial for a host’s responsiveness to environmental change (Zilber-Rosenberg & Rosenberg 2008; Feldhaar 2011; Bordenstein & Theis 2015; Henry *et al*. 2021).

Our study sheds light on the ‘inheritance’ of microbiomes and on hosts as ‘environments’ for the microbiota. While many animals, including a range of insects, have transovarial vertical transmission of bacterial symbionts, i.e. via the egg to the embryo (McFall-Ngai *et al*. 2013; Robinson *et al*. 2019), the microbiome of captive *D. melanogaster*, especially the gut bacteria, are mainly horizontally acquired (Wong *et al*. 2016), and consist mostly of microbes from the diet and from microbes expelled into the immediate environment by conspecifics or predecessors (Blum *et al*. 2013). Interestingly, our results show that the ‘high genetic variation’ bottlenecked populations had a markedly reduced diversity of microbes, low numbers of co-varying bacteria, and lower fitness compared to the outbred control group, despite there being no difference in genetic variation between the two groups (see Ørsted *et al*. 2019 for an elaborated discussion of sampling of alleles during experimental bottlenecks). This could indicate that a reduction in host population size during a population bottleneck constrains the amount of microbial diversity available for random ‘sampling’ for the next generation, similar to the effects of genetic drift on host genetic variation. Indeed, the larger between-line variation in the microbiome community membership, i.e. unweighted UniFrac-based ordination, among the inbred flies as compared to the outbred group strongly suggests a community shift driven by repeated population bottlenecks (**Fig. 3**). Similar to drift, these effects could be exacerbated in small populations, but unlike genetic drift, which is a stochastic process, the effects of bottlenecks yield seemingly directional and predictable effects on microbiome diversity.

The trend of decreasing microbiome diversity with increasing host inbreeding has also been found in inbred populations of Diannan small-ear pigs and tortoises (Yuan *et al*. 2015; Wei *et al*. 2020). This supports that a decreased ability of inbred hosts to harbor diverse and stable microbiomes could be a general pattern, although these studies did not link microbial diversity and genetic variation to fitness effects or evolutionary responses. Previous experiments on *Drosophila* have highlighted the impact of fly and larval density on microbial communities and how this can impact fitness, producing a type of Allee effect (Wertheim *et al*. 2002). Simultaneously, a reduction in fitness due to the genetic effects of population bottlenecks could in turn result in flies that cannot harbor some components of the bacterial community, which could be especially critical if microbes with specific metabolic functions are lost (West *et al*. 2019). We are unable to tease apart these effects in the current study.

Despite this, we show that while host genetic variation has the strongest association with fitness, the microbiome diversity also contributes synergistically to host effects. We envision that increased homozygosity in hosts in a population provides a less optimal habitat for maintaining stable and diverse microbial communities. In this respect, a number of studies have previously reported that hosts which are unhealthy and/or ill harbor different and less diverse microbiomes compared to their healthier counterparts (Mistry *et al*. 2017; Gould *et al*. 2018; Fei *et al*. 2019; Saleem *et al*. 2019; Wagener *et al*. 2020). Moreover, host genotypes have been found to profoundly affect the microbiome composition, as well as their stability and beneficial effects on the host (Bonder *et al*. 2016; Rayes *et al*. 2016; Adair *et al*. 2020), while other studies have shown that microbiomes affect host fitness, development, and even allele frequency and host evolution (Goulet 2015; Wang *et al*. 2018; Rudman *et al*. 2019; Kristensen *et al*. 2021). Given that flies from inbred lines with high nucleotide diversity in our study harbored less diverse microbiomes and had lower fitness than flies from outbred lines which had similar nucleotide diversity, we suggest that the microbiome is important in reducing fly fitness at high levels of genetic variation typical in natural populations. On the other hand, lower nucleotide diversity correlated with even lower microbial diversity and host fitness, suggesting that, when inbreeding results in low host genetic variation, microbial diversity loss is primarily caused by reduced host fitness. Perhaps the less optimal host ‘habitat’ for microbes in lines with low genetic diversity results in less favorable growth conditions. It has been known for a long time that productivity of *Drosophila* can increase when there is high genetic heterogeneity in cultures, presumably because multiple genotypes can better utilize different aspects of the environment (Pérez-Tomé & Toro 1982; Hoffmann & Nielsen 1985). The genetically diverse host populations could provide more variable environments for microbes, allowing a greater diversity of microbes to persist, or support greater microbiome diversity through a higher number of functionally important host genotype-microbial taxon-specific interactions. Tests of these ideas in the context of small populations with restricted genetic variation require further studies that might examine the impact of e.g. antibiotics on host and microbe associations with fitness. Other possibilities for future research in this area involve the transfer of beneficial microbiomes to see if these can “rescue” low fitness populations or individuals suffering from disease. Such experiments have been suggested in other contexts to improve robustness and productivity of livestock and agricultural cultivars and for treating human diseases (Mueller & Sachs 2015; Ser *et al*. 2021). It remains unexplored whether such fitness improvements by microbiome transfer can alleviate the negative impacts commonly associated with low genetic variation like inbreeding depression.

The outbred lines investigated in our study have a much more complex network of co-varying, and interacting microbiomes compared to the low and high genetic variation groups (**Fig. 6**). This suggests that outbred lines harbour healthier and more robust microbial ‘ecosystems’ than bottlenecked lines, where functional redundancy within the diverse microbial community of outbred flies promotes community stability and subsequent host health. This means that removal of individual microbial species is expected to have less impact compared to in low and high genetic variation lines, where the removal of one key species may cause more dramatic effects. The importance of functional redundancy in host-associated microbiome stability and host health has been previously demonstrated in various hosts as well as many other ecosystems (Moya & Ferrer 2016; Liang *et al*. 2020). Notably, it is now widely acknowledged that taxonomically diverse microbiomes harbor robust and stable functional redundancy, where disturbance in the environment is countered, up to a point, by the resilience built upon functional redundancy and high taxonomic diversity in a community (Moya & Ferrer 2016; Liang *et al*. 2020). The global co-variation network in our flies suggests that microbial diversity shifts gradually across a wide continuum of host genetic variation. However, co-variation results from individual fly lines with different levels of genetic diversity more clearly show that the tipping point between community resilience and functional redundancy and communities suffering from more stochastic microbe-microbe associations, and hence reduced functional redundancy, relates more to bottleneck treatment effects than genetic variation within bottleneck treatment. This interpretation is affected by whether lines with high relative abundance of *Wolbachia* was included or not, as *Wolbachia* can affect both host fitness and the presence/abundances of other microbial taxa (see **Supplementary Fig. S9**; Simhadri *et al*. 2017; Audsley *et al*. 2018; Duan *et al*. 2020; Wilches *et al*. 2021).

This bears some resemblance to ecological patterns observed in microbiomes of hosts living in disturbed habitats (Amato *et al*. 2013; Barelli *et al*. 2015; Ingala *et al*. 2019). For instance, howler monkeys inhabiting degraded, more homogenous forests had a less diverse gut microbiome compared to conspecifics in non-fragmented forests, and as a result had lost the microbial metabolic pathway to detoxify plant compounds in the leaves of their diet (Barelli *et al*. 2015). In the same way in which diversity confers resilience in macro-ecological systems (Goodman 1975; Memmott *et al*. 2004), where species-rich communities are less susceptible to invasion as more niches are occupied and limiting resources are used more efficiently, such processes could be important for the robustness of microbial ecosystems within hosts (Lozupone *et al*. 2012; Coyte *et al*. 2015). A high diversity of commensal microbes could mean functional redundancy (Hauffe & Barelli 2019; West *et al*. 2019), thus an increased resilience towards shifts in functional diversity, and better protection against pathogens (Jaenike *et al*. 2010; Koskella & Bergelson 2020).

In conclusion, we demonstrate that restricting host population sizes is associated with a reduction in diversity of host-associated microbiomes. We observe effects of both host genetic variation and microbial diversity in explaining host fitness, however the patterns of causality remain unclear, i.e. we suggest that it is not an all-or-nothing effect, but rather a continuum across nucleotide diversity where fitness and microbiome depletion become relatively more important drivers. It is similarly difficult to establish whether fitness differences are due to beneficial or potential pathogenic bacteria, partly because pathogenicity depends on many factors including composition of the whole microbial community present (Simhadri *et al*. 2017), and age of the host (Ayyaz & Jasper 2013). Despite this, we show clear effects of host population bottlenecks on the diversity of the microbiome, similar to effects normally observed in sampling of genetic alleles during genetic drift, and we show that both the reduction in microbiome diversity and genetic variation is tightly linked to host fitness and evolutionary potential. Therefore, microbial diversity in environmental samples could be a useful proxy for population fitness and potentially adaptability. Lastly, our results open up multiple avenues for further studies, such as transplantation of microbiomes as a means of ‘rescuing’ populations that suffer from inbreeding depression, potentially relevant to species of conservation concern.

## Materials and methods

### Fly population bottlenecks

The *D. melanogaster* flies used in the study originated from 232 wild females caught at Oakridge winery in the Yarra Valley, Victoria, Australia. These females contributed equally to a mass bred population kept at a population size of approximately 1000 individuals. The flies were maintained on a 12:12 h light:dark cycle at 19°C on a standard laboratory food composed of yeast, sucrose, oatmeal, and agar mixed with tap water. Nipagen and acetic acid was added to prevent fungal growth (**Supplementary Table S4**). From the mass bred population, we created 40 lines of each of three different expected levels of inbreeding for a total of 120 lines, from which we could obtain nucleotide diversity measures for 109 lines. This is described in detail in Ørsted *et al*. (2019). In summary, lines from the three levels of inbreeding experienced 2, 3, and 5 consecutive generations of bottlenecks each consisting of two males and two females (census size, N=4) revealing expected inbreeding coefficients of F=0.125, 0.219 and 0.381, respectively. After having reached the desired inbreeding level, we flushed the population sizes to 200 individuals. Simultaneously, we established 10 control lines that were assumed outbred and maintained at minimum 500 individuals.

### Fly lines selected for microbiome analysis

We obtained a measure of the realised genetic variation in all inbred and outbred lines using genotyping-by-sequencing (GBS) to calculate nucleotide diversity (π) from genomic SNPs (described in detail in Ørsted *et al*. (2019); average number of SNPs ± sd = 26,877 ± 4,061). For the microbiome analysis, we aimed at comparing two groups of fly lines: one group with low genetic variation and overall low performance and one with high genetic variation and high overall performance. For a quantitative selection of these groups, we calculated a composite measure of performance as the sum of standardised values of viability and nucleotide diversity (Z-score). We selected the 25 lowest ranking lines and the 25 highest ranking lines based on a summed Z-score (50 lines total; **Supplementary Fig. S1**). Because we selected lines regardless of their inbred/outbred status, nine of the ten outbred control lines were included in the ‘high genetic variation’ lines due to their high Z-scores sum. In all analysis, we differentiate between three groups of flies: ‘low’ and ‘high’ genetic variation lines within the inbred lines and ‘outbred’ (OB) lines that have not been through population bottlenecks. Following establishing the lines, population sizes were flushed to ca. 200 individuals per line, and in the F1 generation egg-to-adult viability was assessed by distributing 15 eggs into each of five vials per line and calculating the proportion of eggs developing successfully to the adult stage (Ørsted *et al*. 2019). Flies used for microbiome analysis consisted of male flies emerging from the egg-to-adult viability assessment (merged across the 5 vials) that were snap frozen in liquid nitrogen at 2-3 days of age and subsequently stored at −80°C.

### 16S rRNA gene sample preparation and sequencing

The whole fly genomic DNA was extracted following the same protocol as has been described in Ørsted *et al*. (2019). In brief, 15 randomly collected male flies from each experimental line was homogenized by bead beating at 2×6 s cycles and 6500 rpm using a Precellys mechanical homogenizer (Bertin Techologies, Montigny le Bretonneux, France), and the DNA was purified with the DNeasy Blood & Tissue kit (QIAGEN, Hilden, Germany) with modifications specific for extracting insect tissues. The V1-V3 hypervariable regions of the bacterial 16S rRNA gene was amplified and multiplexed for Illumina sequencing according to Albertsen *et al*. (2015), and the library pool was sequenced on a MiSeq sequencer using the MiSeq Reagent kit v3, 2×300bp paired-end configuration, and 20% PhiX control spike-in.

### Microbiome analysis

The demultiplexed paired-end reads were quality filtered, assembled to make consensus amplicon reads, clustered into ASVs, and taxonomically assigned using a custom workflow AmpProc version 5.1.0.beta2.11.1 (https://github.com/eyashiro/AmpProc), which relies on USEARCH version 11.0.66_i86linux64 (Edgar 2010) sequenced reads processing and FastTree version 2.1.10 (Price *et al*. 2009) for tree building. When present in insects, the endosymbiont *Wolbachia* can affect host fitness and/or the presence/abundance of other microbial taxa, complicating interpretation of results (Simhadri *et al*. 2017; Audsley *et al*. 2018; Duan *et al*. 2020; Wilches *et al*. 2021). Therefore, we removed six lines with a relative *Wolbachia* abundance >85 %. These six lines were evenly distributed among the genetic variation groups (**Supplementary Fig. S8**). This threshold was chosen to maintain a relatively high number of reads per line/sample, i.e., at least 14,000 reads per sample, after *Wolbachia* reads were removed. Prior to further analysis, we also tested whether removing *Wolbachia* from the frequency table would skew our results and interpretations. There was no difference in relative abundance of *Wolbachia* between genetic variation groups (Kruskal-Wallis Rank Sum test; χ^2^ = 2.54, df = 2, *p* = 0.28). Further, we found no significant correlations between *Wolbachia* abundance and nucleotide diversity (r_s_ = 0.24, t_48_ = 1.68, *p* = 0.10) or fitness (r_s_ = 0.23, t_48_ = 1.67, *p* = 0.10), and there was no significant difference between the six removed lines and the 44 remaining lines in mean nucleotide diversity (Wilcoxon rank sum test; W = 110, *p* = 0.53) or mean fitness (W = 116, *p* = 0.64). Thus, our main results namely that lines with low genetic variation and fitness harbor a less diverse microbiome were not affected by removing lines high in *Wolbachia* abundance. However, the prevalence of *Wolbachia* reads above 10% per sample distorted the relative abundance of other bacterial taxonomic groups to a small extent (**Supplementary Fig. S8D**), thereby leading to slightly different interpretations of the co-abundance networks (see **Supplementary Fig S9** for details).

Next, QIIME version 1.9.1 (Caporaso *et al*. 2010) was used to rarefy all of the samples to the smallest acceptable sample size i.e. 14,488 reads per sample (single_rarefaction.py), and to generate the observed and estimated chao1 richness and Shannon-Wiener and Simpsons diversity values for each sample (alpha_diversity.py), weighted and unweighted UniFrac matrices (beta_diversity.py), and core microbiome groups (compute_core_microbiome.py). R version 3.5.0 was used for downstream analysis and statistics. To assess the composition of core microbiomes, we first normalized the dataset to eight lines per fly group (which is the number of outbred lines in the study after filtering samples with low read counts and removing *Wolbachia*) by randomly keeping three, two, and three lines from the high, medium, and low inbreeding levels, respectively, from the high and low genetic variation groups (Ørsted *et al*. 2019), and all of the outbred lines. Next, we calculated as belonging to the core microbiome, the percentage to the nearest tenth if we were to accept a prevalence of an ASV in 80% of the lines in each fly group.

Non-metric multidimensional plots based on the UniFrac matrices were generated with the R-package ‘vegan’ (version 2.5-2 (Oksanen *et al*. 2018)). The effect of two host traits (nucleotide diversity, egg-to-adult viability; Ørsted *et al*. 2019) were calculated with the envfit function and the significant envfit values (*p* < 0.05) of the fly host traits were displayed as arrows and fly host trait gradients across the ordination space as isolines using the ordisurf function. PERMANOVA analysis using vegan’s adonis function was undertaken for the UniFrac matrices as a function of fly lines, nucleotide diversity and viability. Differentially enriched ASVs were identified with DESeq2 (version 1.22.2 (Love *et al*. 2014)) using the Wald test, contrasting across three levels (low and high genetic variation and outbred fly groups), and accepting the enriched ASVs with adjusted *p* < 0.05. The same approach was used on the genus-level frequency table. To test for differences in measures of microbiome diversity between the genetic variation groups, we performed Kruskal-Wallis tests and pairwise Wilcox’s t-tests between groups.

### Generalised linear mixed models

To assess drivers of fitness measured as egg-to-adult viability, we fitted generalised linear mixed effect models (GLMMs) in the R-package ‘lme4’ (Bates *et al*. 2015) assuming a binomial distribution with logit link function. We fitted viability as a function of nucleotide diversity and microbiome diversity (four different measures as described above) and their interaction as fixed effects; replicate vials were included as a random effect, as flies from the same vial are not independent. The full models were compared with individual univariate models of either nucleotide diversity or microbiome diversity by χ^2^ difference tests. We detected no over-dispersion in any of the models (residuals to df ratio > 0.31; χ^2^ > 76.02; *p* > 0.99). Conditional coefficients of determination of the GLMMs interpreted as the variance explained by the entire model, including both fixed and random effects, were calculated as 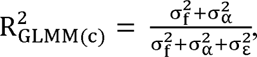 where 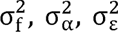 are the variances of the fixed effect components, the random effects and the residual variance, respectively (see ‘delta-method’ in Nakagawa *et al*. (2017)).

### Co-occurrence network analysis

For co-occurrence network analysis, the ASV table of all of the fly lines was used for the global network, and the subset of ASV tables generated for the core microbiomes analysis for outbred, high genetic variation, and low genetic variation groups were used as the starting frequency tables. In each of these ASV frequency tables, the fly nucleotide diversity and egg-to-adult viability measurements were included. The ASVs present in fewer than three lines and ASVs with less than 30 reads in the respective frequency tables were removed. The ‘Hmisc’ R-package (Harrell 2021) was used to generate Spearman correlation matrices, and the upper triangle was used to generate the network edge list. Only edges with FDR-corrected *p* < 0.05 and correlation coefficients > 0.5 (or < −0.5) were kept. In Gephi version 0.9.2, the network was generated with the Fruchterman Reingold algorithm and minor crossing of edges were corrected manually to improve the visualization of the clusters. From among Gephi’s built-in features, the Leiden algorithm was used to calculate the modularity structure using edge weights and default settings, and the degree of connectivity was calculated as the number of nodes with which each node is connected by an edge.

## Acknowledgements

We thank Kelly Richardson for providing the fly stock population used in the experiments, and Kelly and Perran Stott-Ross for their help in setting up the inbred lines. We thank Kåre L. Nielsen, and Elsa Sverrisdóttir for bioinformatics assistance, and Nancy Endersby-Harshman, Helle Blendstrup, Susan M. Hansen, and Neda N. Moghadam for laboratory assistance.

## Supplementary Information

**Supplementary Fig. S1.**
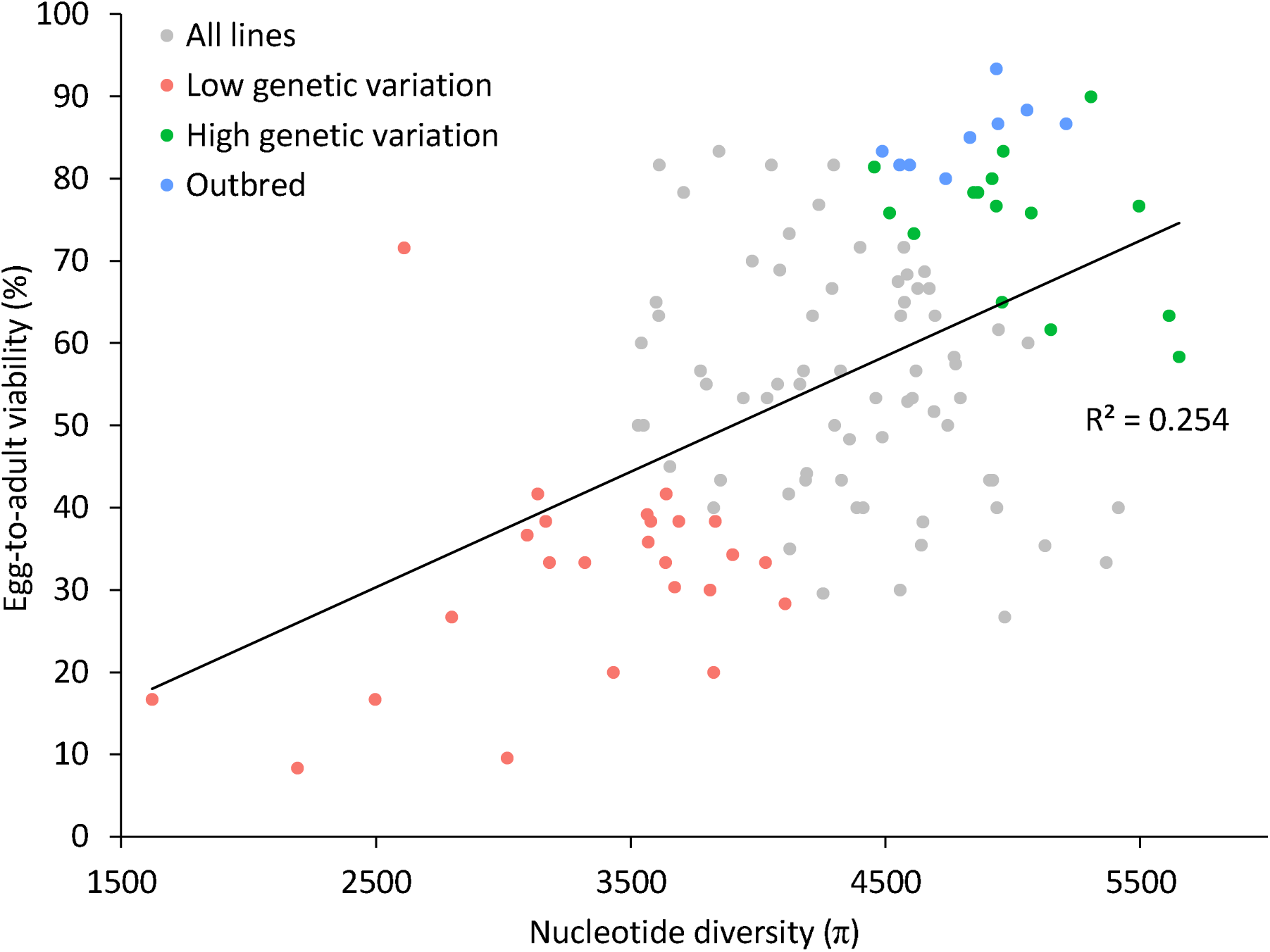
Scatterplot of line mean egg-to-adult viability (%) and nucleotide diversity (π) as obtained through GBS of 119 lines (grey circles; from Ørsted et al. 2019). From these lines, we selected three groups of lines for analysis of the microbiome, 25 lines with low overall performance (red circles), and 25 lines with high performance (green and blue). This selection was based on a composite measure of the overall performance calculated as the sum of standardized viability and nucleotide diversity. Because we selected lines regardless of their inbred/outbred status, the ‘high’ group included nine of the outbred controls as they generally had very high fitness and genetic variation. To distinguish between the effects of genetic variation within inbred lines and population bottlenecks, we therefore differentiate between three genetic variation categories: ‘Low genetic variation’ and ‘High genetic variation’ lines and ‘Outbred’ (OB) control lines. The solid line represents the linear regression on all lines with the R^2^ value shown.

**Supplementary Fig. S2.**
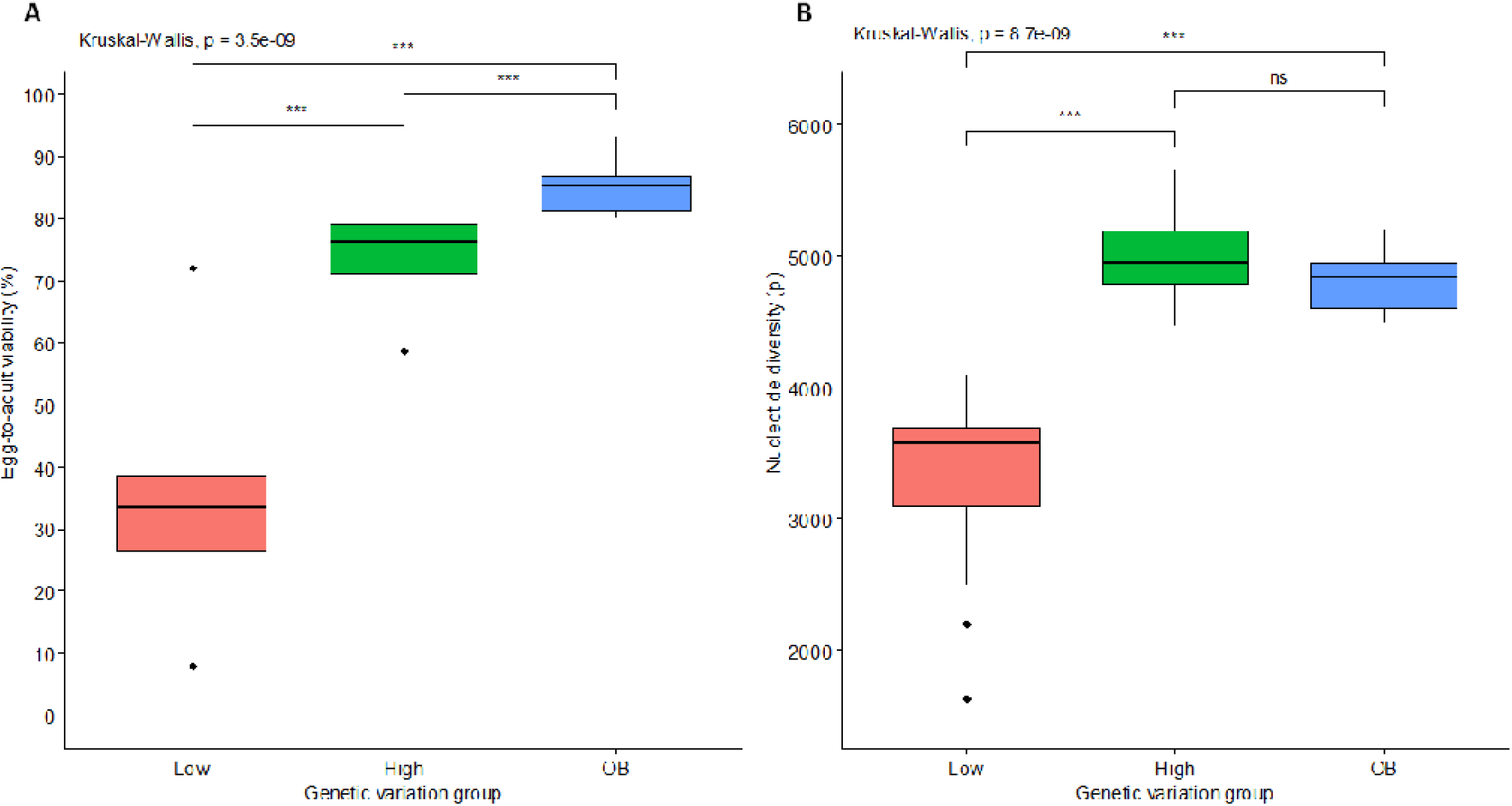
Both the high genetic variation (‘High’) and the outbred (‘OB’) lines exhibited a significantly higher viability (**A**) and nucleotide diversity (**B**) than the low genetic variation lines (‘Low’), and viability of OB lines was higher than that of the high genetic variation lines, while nucleotide diversity was indistinguishable between these two groups (two-sample Wilcox’s t-tests; *p* < 0.05). The *p* values of a Kruskal-Wallis test show significant effects of group, while asterisks denote the results of pairwise Wilcoxon’s t-tests between groups; *** *p* < 0.001; ** *p* < 0.01; * *p* < 0.05; and ns: *p* > 0.05.

**Supplementary Fig. S3.**
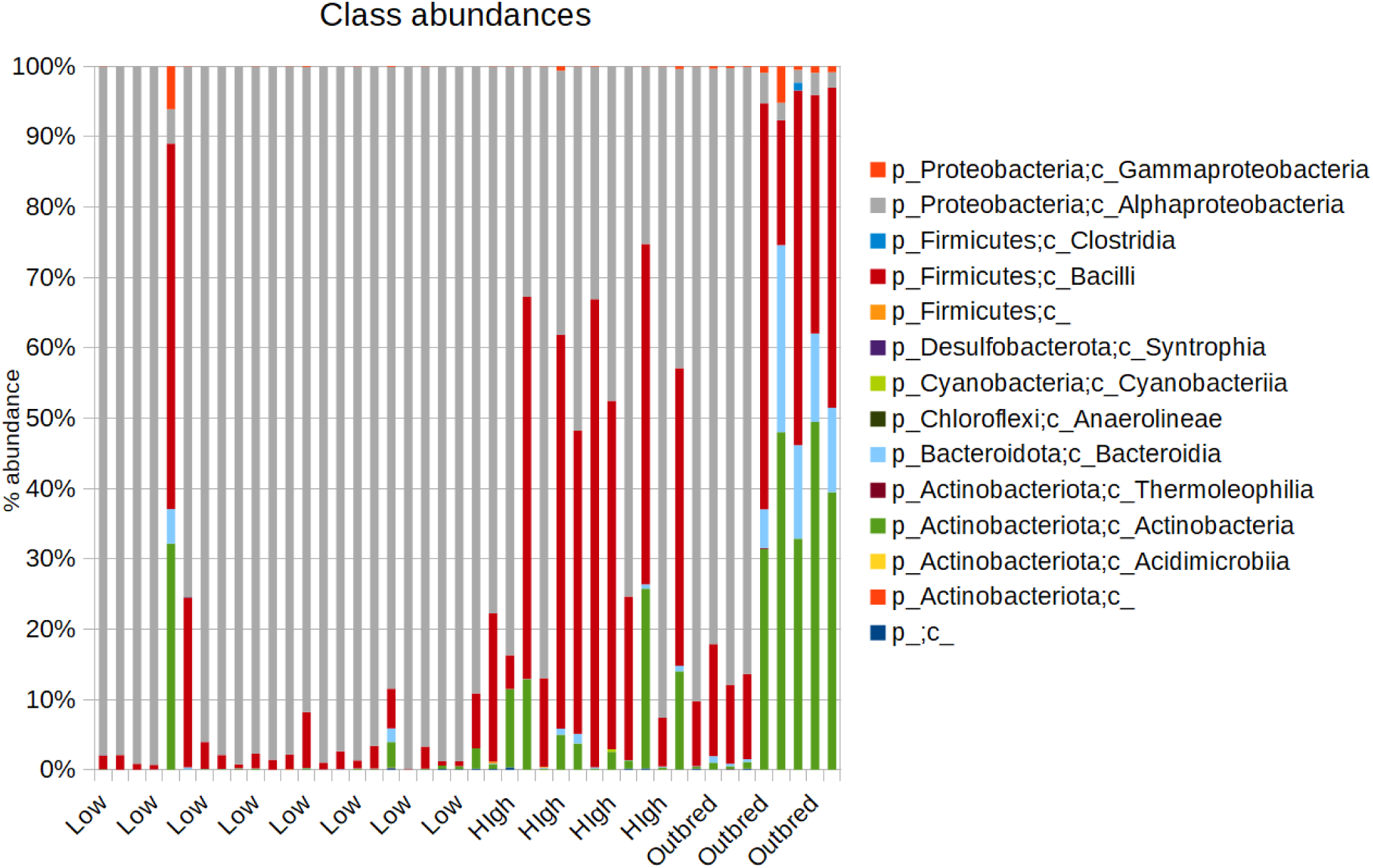
Bar chart of class-level taxonomic composition of the fly microbiomes in the three major groups of host genetic variation (Low: low genetic variation, High: high genetic variation, and Outbred: outbred flies).

**Supplementary Fig. S4.**
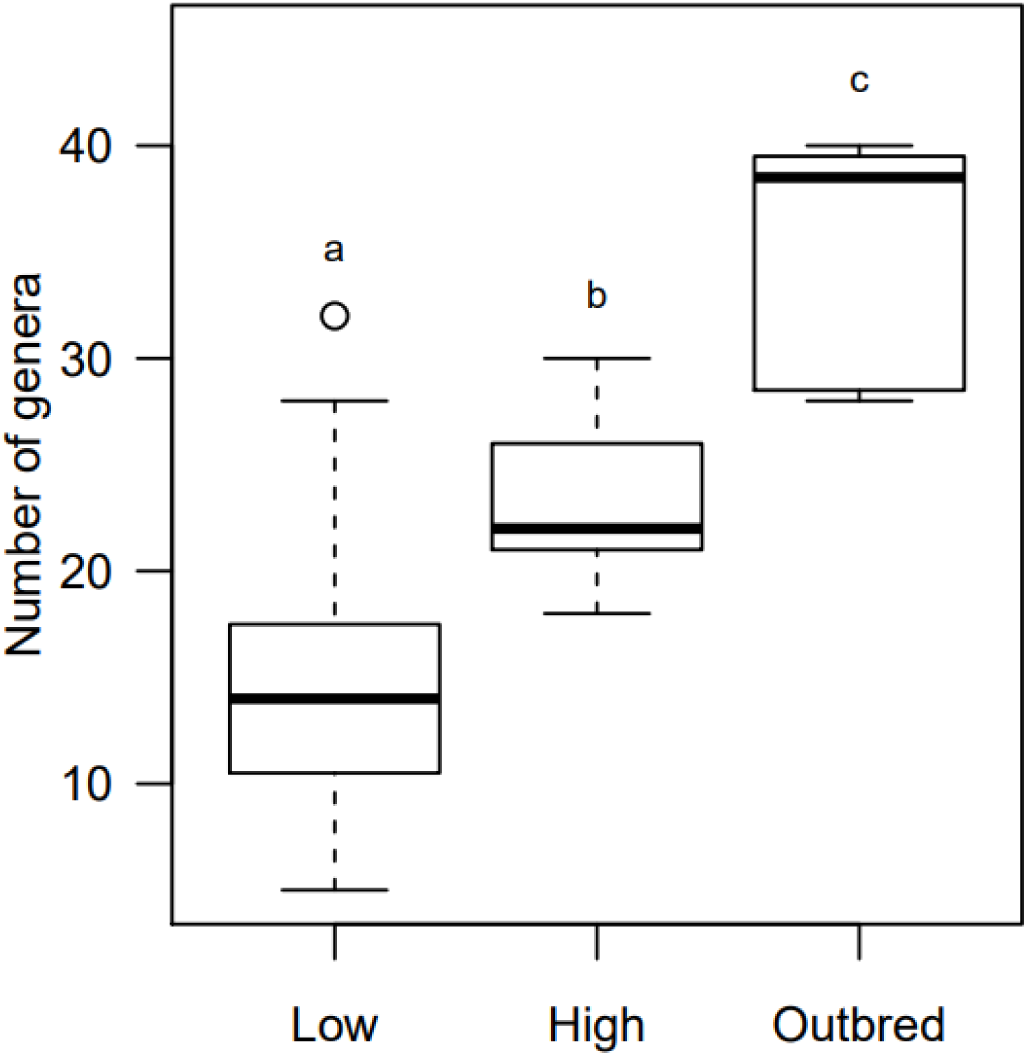
Box plot of the number of bacterial genera harboured in each line, grouped according to fitness (Low: low genetic variation, High: high genetic variation, and Outbred: outbred flies). Letters denote significant differences in mean number of genera (Tukey’s HSD test, *p* < 0.05).

**Supplementary Fig. S5.**
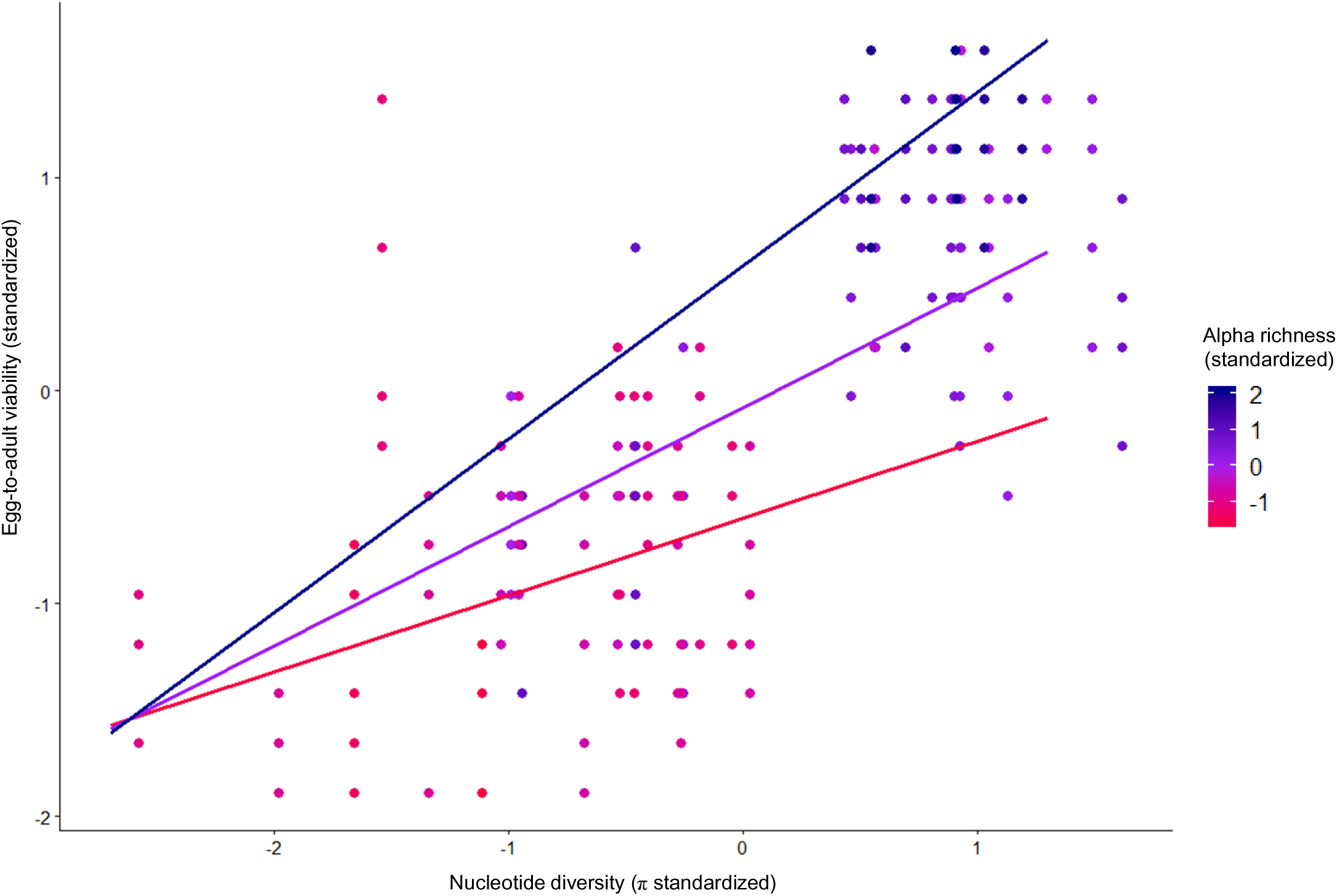
Scatterplot of egg-to-adult viability (z-standardised) from 5 vials per line as a function of nucleotide diversity (π z-standardised) for each of the 44 lines investigated for microbial diversity; the colour of points denote observed ASV richness (alpha richness, z-standardised). The three regression lines represent regressions on lines with the highest (blue), mean (purple), and lowest (red) alpha richness, visualizing the significant positive interaction between nucleotide diversity and microbiome richness in explaining fitness of the host population (see **Table 1** for GLMM results).

**Supplementary Fig. S6.**
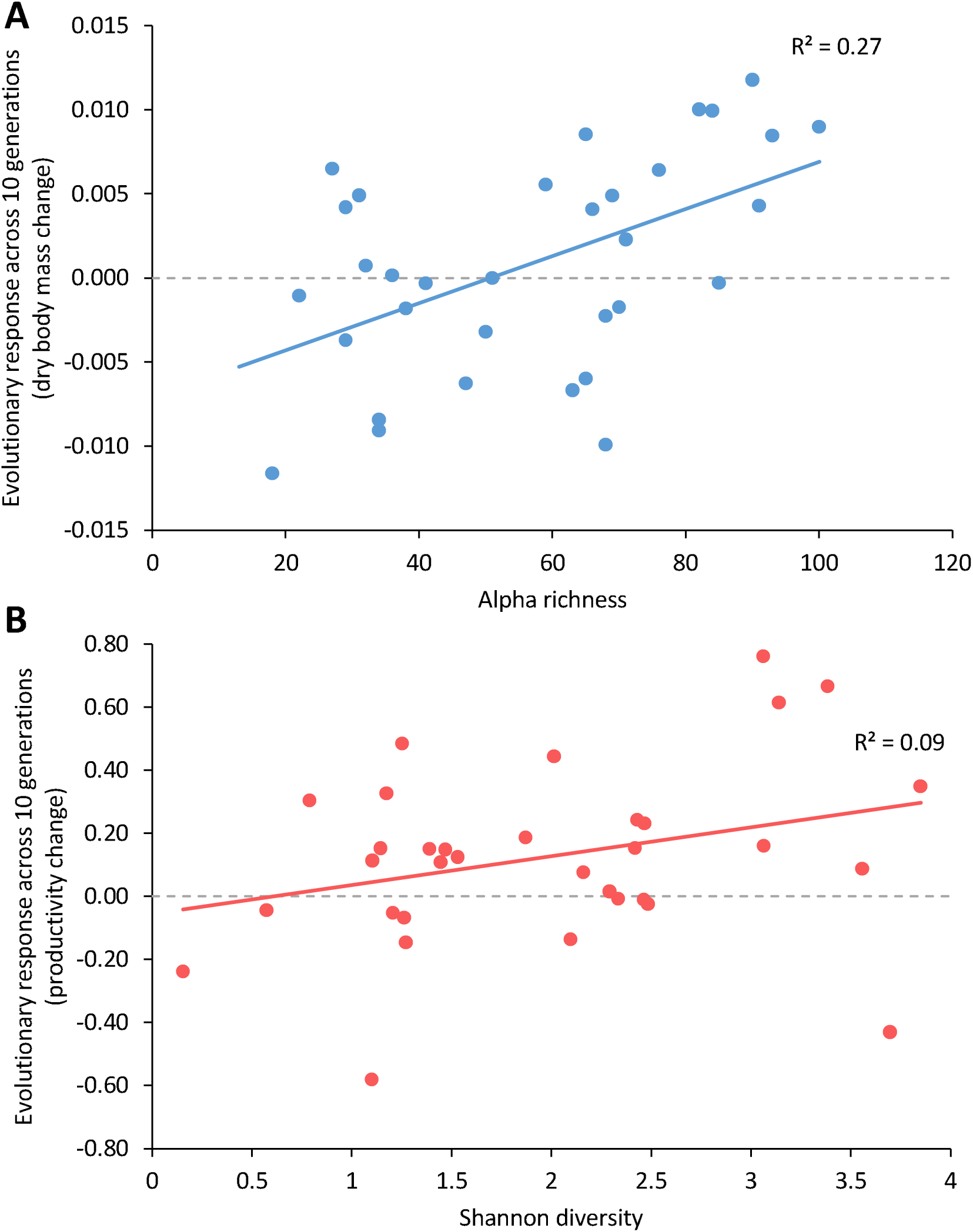
Using data from Ørsted *et al*. (2019) from the set of lines in the present study, we associate microbial diversity with evolutionary responses in two traits: **A.** dry body mass (in mg), and **B.** productivity (in eggs per female per day). Here evolutionary responses are defined as the slope of an ordinary linear regression across 10 generations of rearing on a stressful medium. We correlated all four measures of microbial diversity used in the present study to these two traits (eight comparisons), but here we only included the cases where there was a significant effect of microbial diversity. For both traits, we found an effect of microbial diversity (alpha richness regressed on response in dry body mass: *t*_29_ = 2.09, *p* = 0.045, and Simpson’s diversity regressed on response in productivity: *t*_29_ = 2.42, *p* = 0.022), but no effect of genetic variation, π (*t*_29_ = −1.07, *p* = 0.92, and *t*_29_ = −0.14, *p* = 0.89, for dry body mass and productivity, respectively). For details on assessment of these phenotypes, see Ørsted *et al*. (2019).

**Supplementary Fig. S7.**
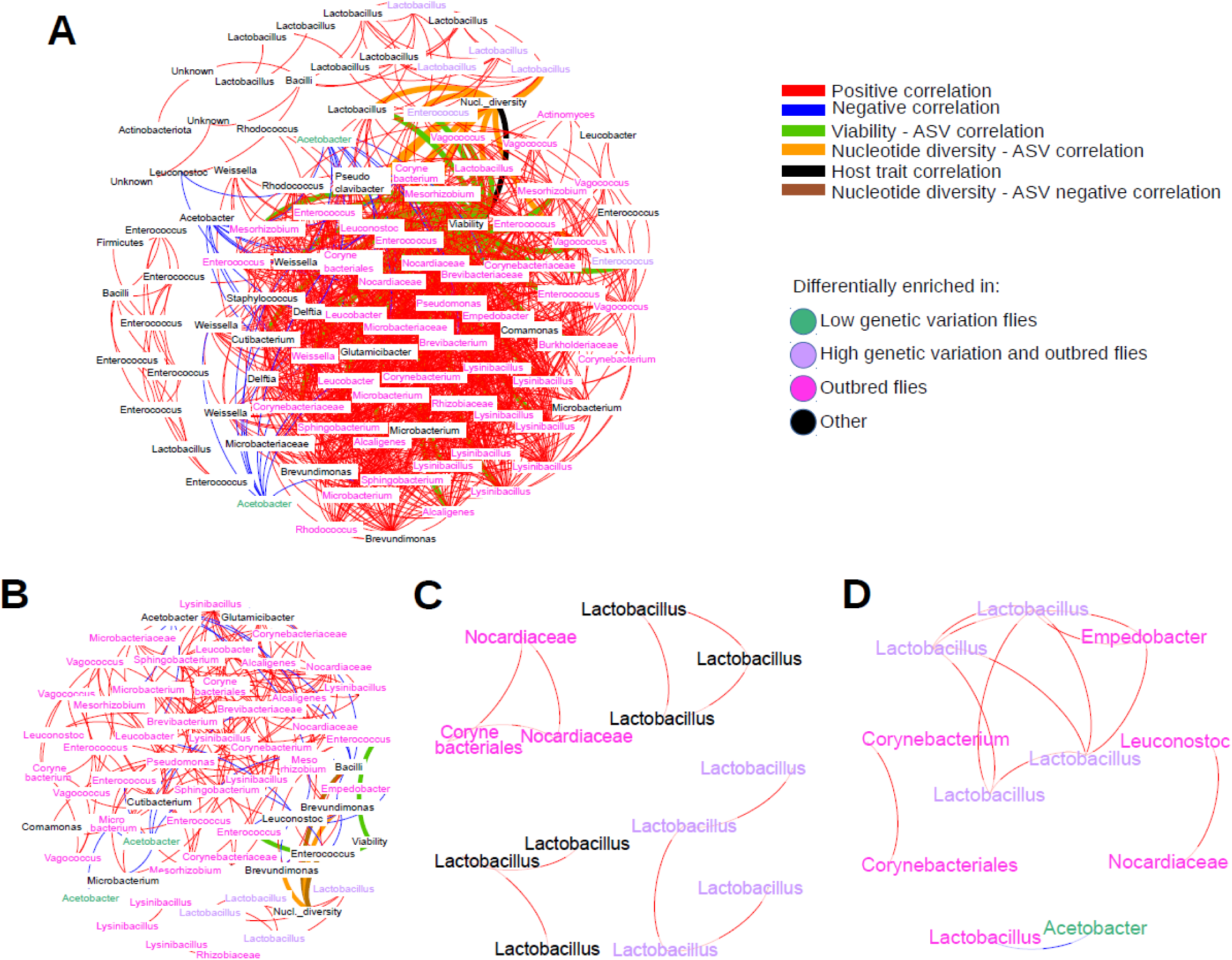
Co-abundance networks of the fly microbiome (closely resembles **Fig. 6** in main text, except here we supply the ASVs for each node). ASVs present in at least 30 reads in total and in at least three lines, and correlations of > 0.5 (or <-0.5) and fdr-corrected *p*-values of < 0.05 are shown. The nodes are individual ASVs and the host fitness traits, egg-to-adult viability (Viability) and nucleotide diversity (Nucl. diversity), while the edges represent positive and negative correlations, and correlations linking host fitness traits and bacterial ASVs (which were positive correlations). The network containing lines from all fly groups (**A**), the outbred (**B**), high genetic variation (**C**), and low genetic variation (**D**) groups are shown with the taxonomic assignment of each node. Non-highlighted nodes that could not be taxonomically assigned are left as ASV identifier numbers. The highlighted nodes are color-coded according to the major DESeq differential abundance groups. The results of both the Leiden algorithm and degree were plotted into the network graphs as node colors and node sizes, respectively. For cross-referencing the DESeq2 differentially enriched ASVs in the networks, only the most robust differential abundance patterns were displayed, notably the ASVs that were systematically enriched in the low genetic variation flies in both the low-high and low-outbred pairwise comparisons (low), the ASVs systematically enriched in the outbred flies (outbred), and those depleted in the low genetic variation flies (*vs.* high & outbred).

**Supplementary Fig. S8.**
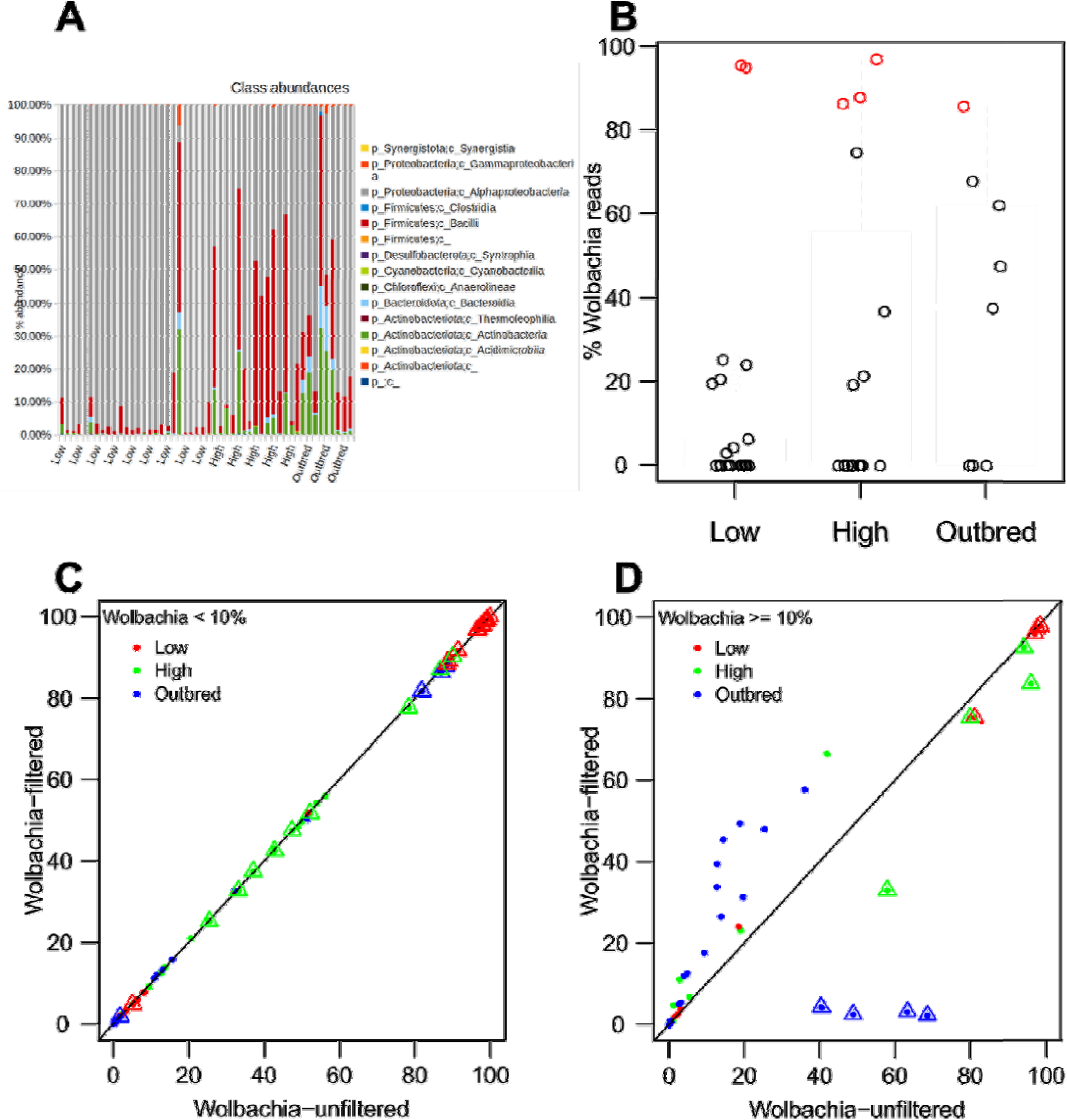
(A) Bar chart of class-level taxonomic composition of the fly microbiomes in the three major groups of host genetic diversity when *Wolbachia* reads are included in the frequency table. (**B**) Relative abundance of the endosymbiont *Wolbachia* ASV, across all 50 initially selected lines . We removed six lines total with abundances above a 85 % threshold (red circles; two low genetic variation, three high genetic variation and one outbred). No effect of genetic variation group on relative abundance was found (Kruskal-Wallis rank sum test; χ^2^ = 2.54, df = 2, *p* = 0.28). (**C**) The percent proportion of the bacterial class-level taxa from the lines containing less than 10% *Wolbachia* were plotted for the dataset where *Wolbachia* was included and for the dataset where *Wolbachia* was removed. (**D**) The proportion of the class-level taxa from lines with 10% or more *Wolbachia* reads were also plotted. Each dot corresponds to one class from one sample. Given that *Wolbachia* belong to the Alphaproteobacteria, the dots corresponding to Alphaproteobacteria are overlaid with triangles. The regression line indicates the plot region where taxon proportions are 1:1 ratio between the two axes. When *Wolbachia* abundance was low, the community profile was highly similar between the dataset with and without *Wolbachia*, whereas, when *Wolbachia* abundance was high, their prevalence skewed the relative proportion of the other taxonomic groups.

**Supplementary Fig. S9.**
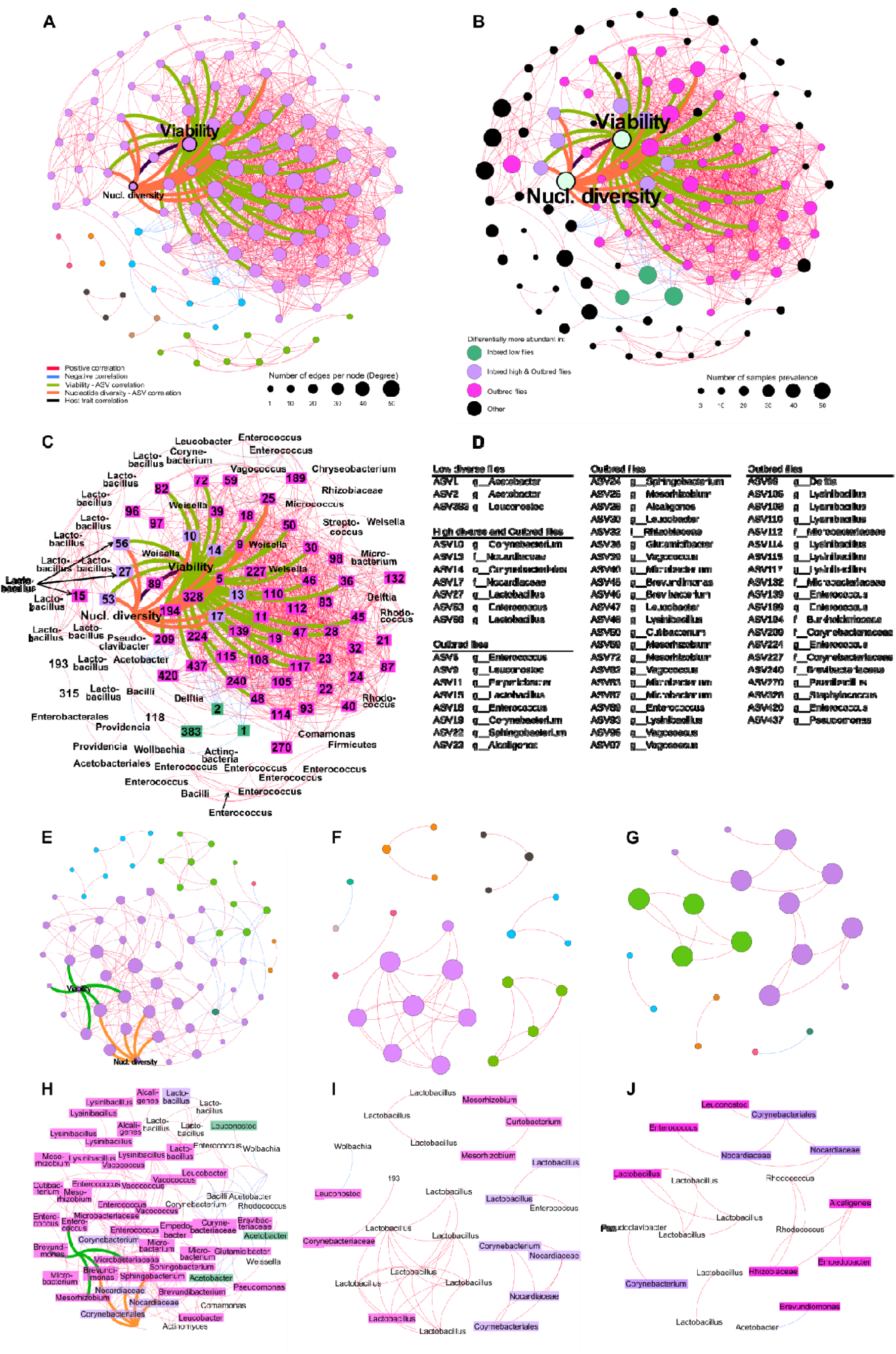
Co-abundance networks of the fly microbiome using the ASV table that includes *Wolbachia*. ASVs present in at least 30 reads in total and in at least three lines; correlations > 0.5 (or <-0.5) and fdr-corrected p-values < 0.05 are shown. The nodes are individual ASVs and the host fitness traits, egg-to-adult viability (Viability) and nucleotide diversity (Nucl. diversity), while the edges represent positive and negative correlations, and correlations linking host fitness traits and bacterial ASVs (which were positive correlations). Panels **A, B, E, F, G** are corresponding networks found in **Fig. 6**, while **C, D, H, I, J** are corresponding networks in **Supplementary Fig. S7.** The numerical values in the coloured nodes in panel **C** correspond to the ASV IDs in panel **D**. The conclusions that can be made from the dataset which included *Wolbachia* differ from conclusions made with the dataset excluding the endosymbiont. Therefore, **Fig. S9 E-G** shows that reducing fly genetic variation gradually decreases microbe-microbe associations in the fly microbiome, while **Fig. 7 C-E** shows that fly genetic variation and microbe-microbe associations do not monotonically decrease; rather, inbred flies host microbes that do not interact much compared to outbred flies.

**Supplementary Table S1**

Table S1 found online in the Supplementary Information as a Microsoft Excel spreadsheet. Nucleotide diversity, fitness and microbiome diversity (observed alpha richness, estimated chao1 richness, Shannon-Wiener and Simpson’s indices), and ASV table rarefied to 14,488 reads per sample of the 44 fly lines used in the main results, as well as ASV table rarefied to 29,000 reads per sample where *Wolbachia* was not prefiltered, and which includes the six lines that were removed from the original 50 lines due to high relative *Wolbachia* abundance (in a separate tab). Headers of the metadata sheets are described in the last tab.

**Supplementary Table S2.** Results of general linear mixed models (GLMMs) of egg-to-adult viability as a function of nucleotide diversity (NuclDiv) and microbiome diversity and their interaction as fixed effects. The measures of microbiome diversity are divided into alpha richness indices; estimated ASV richness (chao1), and diversity indices accounting for relative abundances as well (Shannon-Wiener index (Shannon) and Simpson’s index (Simpson) in each of their own model. Both dependent and independent variables are scaled (Z-standardization) to allow direct comparison of effect sizes. Replicate vial ID were included as a random effect, as flies from the same vial are not considered independent. Conditional coefficients of determination of the GLMMs 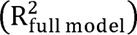 interpreted as the variance explained by the entire model, including both fixed and random effects, is shown. Asterisks denote the significance of individual variables or interactions; *** *p* < 0.001; ** *p* < 0.01; and * *p* < 0.05. The full model including both dependent variables and their interaction is compared with individual models with either nucleotide diversity or alpha diversity with a χ^2^ test.

| Microbiome diversity measure | Fixed effects | Estimate | Std. Error | z value | p |
| --- | --- | --- | --- | --- | --- |
| chao1 estimated ASV Richness (chao1)<br>$R^2_{\text{full model}} = 0.353$ | Intercept | 0.134 | 0.081 | 1.658 | 0.0972 |
|  | NuclDiv | 0.767 | 0.084 | 9.173 | 4.61E-20 *** |
|  | chao1 | 0.490 | 0.083 | 5.882 | 4.04E-09 *** |
|  | NuclDiv*chao1 | 0.141 | 0.086 | 1.632 | 0.1026 |
|  | **Random effects** | **Std.Dev** |  |  |  |
|  | Vial | 0.707 |  |  |  |
| | **Comparison with individual models** | $\chi^2$ df | $\chi^2$ | p | |
|  | NuclDiv; chao1; Full model | 2 | 75.434 | 2.00E-16 | *** |
|  | Fixed effects | Estimate | Std. Error | z value | p |
| Shannon-Wiener Index (Shannon)<br>$R^2_{\text{full model}} = 0.354$ | Intercept | 0.148 | 0.085 | 1.736 | 0.0826 |
|  | NuclDiv | 0.817 | 0.097 | 8.411 | 4.07E-17 *** |
|  | Shannon | 0.411 | 0.093 | 4.430 | 9.41E-06 *** |
|  | NuclDiv*Shannon | 0.105 | 0.088 | 1.193 | 0.2329 |
|  | **Random effects** | **Std.Dev** |  |  |  |
|  | Vial | 0.735 |  |  |  |
| | **Comparison with individual models** | $\chi^2$ df | $\chi^2$ | p | |
|  | NuclDiv; Shannon; Full model | 2 | 67.413 | 2.30E-15 | *** |
|  | Fixed effects | Estimate | Std. Error | z value | p |
| Simpson's Index (Simpson)<br>$R^2_{\text{full model}} = 0.354$ | Intercept | 0.130 | 0.078 | 1.660 | 0.0970 |
|  | NuclDiv | 0.896 | 0.090 | 9.997 | 1.57E-23 *** |
|  | Simpson | 0.341 | 0.085 | 4.023 | 5.74E-05 *** |
|  | NuclDiv*Simpson | 0.137 | 0.073 | 1.879 | 0.0603 |
|  | **Random effects** | **Std.Dev** |  |  |  |
|  | Vial | 0.750957 |  |  |  |
| | **Comparison with individual models** | $\chi^2$ df | $\chi^2$ | p | |
|  | NuclDiv; Simpson; Full model | 2 | 86.078 | 2.00E-16 | *** |

**Supplementary Table S3**

DESeq2 table is found online as a Microsoft Excel spreadsheet. The log fold change calculations are done on the ASV (L7 tabs in the spreadsheet) and genus level frequency tables (L6 tabs in the spreadsheet). The DESeq contrasts were done among outbred, high and low genetic variation fly groups, and the pairwise contrasts are displayed in separate sheets. For the most significant pairwise comparisons (*p* < 0.1), the fly group in which the ASVs or genera are enriched is indicated.

**Supplementary Table S4.**
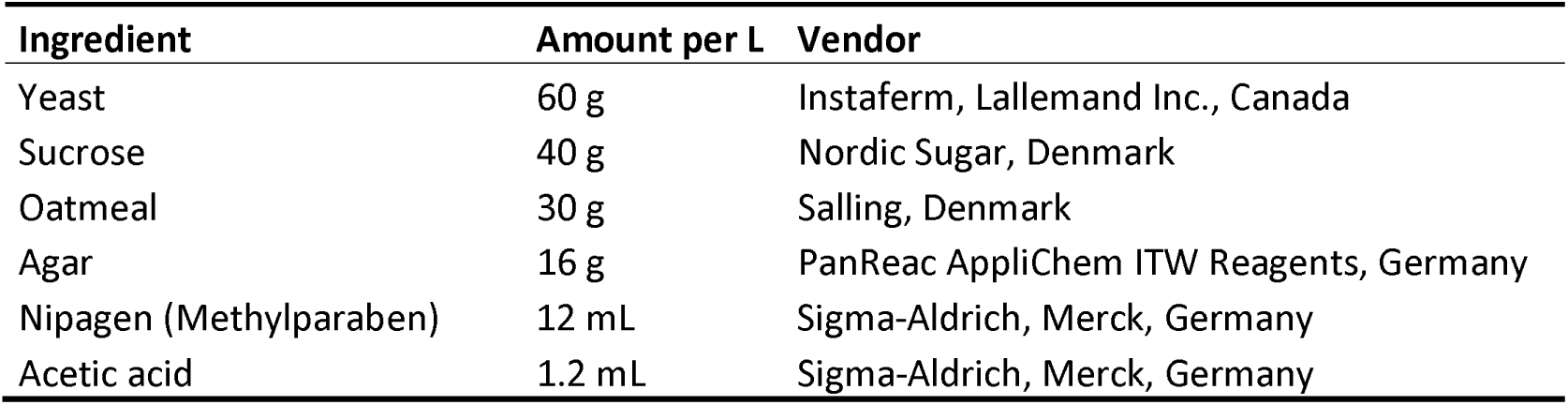
Amount per liter and vendor of standard fly medium ingredients used in the present study

| Ingredient | Amount per L | Vendor |
| --- | --- | --- |
| Yeast | 60 g | Instaferm, Lallemand Inc., Canada |
| Sucrose | 40 g | Nordic Sugar, Denmark |
| Oatmeal | 30 g | Salling, Denmark |
| Agar | 16 g | PanReac AppliChem ITW Reagents, Germany |
| Nipagen (Methylparaben) | 12 mL | Sigma-Aldrich, Merck, Germany |
| Acetic acid | 1.2 mL | Sigma-Aldrich, Merck, Germany |

## References

Adair, K.L., Bost, A., Bueno, E., Kaunisto, S., Kortet, R., Peters-Schulze, G., et al. (2020). Host determinants of among-species variation in microbiome composition in drosophilid flies. The ISME Journal, 14, 217–229.

Albertsen, M., Karst, S.M., Ziegler, A.S., Kirkegaard, R.H. & Nielsen, P.H. (2015). Back to basics – the influence of DNA extraction and primer choice on phylogenetic analysis of activated sludge communities. PLOS ONE, 10, e0132783.

Amato, K.R., Yeoman, C.J., Kent, A., Righini, N., Carbonero, F., Estrada, A., et al. (2013). Habitat degradation impacts black howler monkey (*Alouatta pigra*) gastrointestinal microbiomes. The ISME Journal, 7, 1344–1353.

Audsley, M.D., Seleznev, A., Joubert, D.A., Woolfit, M., O’Neill, S.L. & McGraw, E.A. (2018). *Wolbachia* infection alters the relative abundance of resident bacteria in adult *Aedes aegypti* mosquitoes, but not larvae. Molecular Ecology, 27, 297–309.

Ayyaz, A. & Jasper, H. (2013). Intestinal inflammation and stem cell homeostasis in aging *Drosophila melanogaster*. Frontiers in Cellular and Infection Microbiology, 3, 98.

Bahrndorff, S., Alemu, T., Alemneh, T. & Lund Nielsen, J. (2016). The microbiome of animals: implications for conservation biology. International Journal of Genomics, 2016, 5304028.

Barelli, C., Albanese, D., Donati, C., Pindo, M., Dallago, C., Rovero, F., et al. (2015). Habitat fragmentation is associated to gut microbiota diversity of an endangered primate: implications for conservation. Scientific Reports, 5, 14862.

Bates, D., Mächler, M., Bolker, B. & Walker, S. (2015). Fitting linear mixed-effects models using lme4. Journal of Statistical Software, 67, 1–48.

Blum, J.E., Fischer, C.N., Miles, J. & Handelsman, J. (2013). Frequent replenishment sustains the beneficial microbiome of *Drosophila melanogaster*. mBio, 4, e00860–13.

Bonder, M.J., Kurilshikov, A., Tigchelaar, E.F., Mujagic, Z., Imhann, F., Vila, A.V., et al. (2016). The effect of host genetics on the gut microbiome. Nature Genetics, 48, 1407–1412.

Bordenstein, S.R. & Theis, K.R. (2015). Host biology in light of the microbiome: ten principles of holobionts and hologenomes. PLOS Biology, 13, e1002226.

von Burg, S., Ferrari, J., Müller, C.B. & Vorburger, C. (2008). Genetic variation and covariation of susceptibility to parasitoids in the aphid *Myzus persicae*: no evidence for trade-offs. Proceedings of the Royal Society B: Biological Sciences, 275, 1089–1094.

Caporaso, J.G., Kuczynski, J., Stombaugh, J., Bittinger, K., Bushman, F.D., Costello, E.K., et al. (2010). QIIME allows analysis of high-throughput community sequencing data. Nature methods, 7, 335–336.

Chandler, J.A., Morgan Lang, J., Bhatnagar, S., Eisen, J.A. & Kopp, A. (2011). Bacterial communities of diverse *Drosophila* species: ecological context of a host–microbe model system. PLOS Genetics, 7, e1002272.

Chaston, J.M., Dobson, A.J., Newell, P.D. & Douglas, A.E. (2016). Host genetic control of the microbiota mediates the *Drosophila* nutritional phenotype. Applied and Environmental Microbiology, 82, 671–679.

Chong, R.A. & Moran, N.A. (2016). Intraspecific genetic variation in hosts affects regulation of obligate heritable symbionts. Proceedings of the National Academy of Sciences, 113, 13114– 13119.

Coyte, K.Z., Schluter, J. & Foster, K.R. (2015). The ecology of the microbiome: networks, competition, and stability. Science, 350, 663–666.

Cullen, C.M., Aneja, K.K., Beyhan, S., Cho, C.E., Woloszynek, S., Convertino, M., et al. (2020). Emerging priorities for microbiome research. Frontiers in Microbiology, 11, 136.

Dobson, A.J., Chaston, J.M., Newell, P.D., Donahue, L., Hermann, S.L., Sannino, D.R., et al. (2015). Host genetic determinants of microbiota-dependent nutrition revealed by genome-wide analysis of Drosophila melanogaster. Nature Communications, 6, 6312.

Douglas, A.E. (2019). Simple animal models for microbiome research. Nature Reviews Microbiology, 17, 764–775.

Du, E.J., Ahn, T.J., Kwon, I., Lee, J.H., Park, J.-H., Park, S.H., et al. (2016). TrpA1 regulates defecation of food-borne pathogens under the control of the Duox pathway. PLOS Genetics, 12, e1005773.

Duan, X.-Z., Sun, J.-T., Wang, L.-T., Shu, X.-H., Guo, Y., Keiichiro, M., et al. (2020). Recent infection by *Wolbachia* alters microbial communities in wild *Laodelphax striatellus* populations. Microbiome, 8, 104.

Edgar, R.C. (2010). Search and clustering orders of magnitude faster than BLAST. Bioinformatics, 26, 2460–2461.

Engelstädter, J. & Hurst, G.D.D. (2009). The ecology and evolution of microbes that manipulate host reproduction. Annual Review of Ecology, Evolution, and Systematics, 40, 127–149.

Ericsson, A.C. & Franklin, C.L. (2015). Manipulating the gut microbiota: methods and challenges. ILAR Journal, 56, 205–217.

Fei, N., Bernabé, B.P., Lie, L., Baghdan, D., Bedu-Addo, K., Plange-Rhule, J., et al. (2019). The human microbiota is associated with cardiometabolic risk across the epidemiologic transition. PLOS ONE, 14, e0215262.

Feldhaar, H. (2011). Bacterial symbionts as mediators of ecologically important traits of insect hosts. Ecological Entomology, 36, 533–543.

Ferrari, J. & Vavre, F. (2011). Bacterial symbionts in insects or the story of communities affecting communities. Philosophical Transactions of the Royal Society B: Biological Sciences, 366, 1389–1400.

Frankham, R. (2005). Genetics and extinction. Biological Conservation, 126, 131–140.

Goodacre, S.L., Martin, O.Y., Bonte, D., Hutchings, L., Woolley, C., Ibrahim, K., et al. (2009). Microbial modification of host long-distance dispersal capacity. BMC Biology, 7, 32.

Goodman, D. (1975). The theory of diversity-stability relationships in ecology. The Quarterly Review of Biology, 50, 237–266.

Gould, A.L., Zhang, V., Lamberti, L., Jones, E.W., Obadia, B., Korasidis, N., et al. (2018). Microbiome interactions shape host fitness. Proceedings of the National Academy of Sciences, 115, E11951–E11960.

Goulet, O. (2015). Potential role of the intestinal microbiota in programming health and disease. Nutrition Reviews, 73, 32–40.

Grueber, C.E., Peel, E., Wright, B., Hogg, C.J., Belov, K., Grueber, C.E., et al. (2019). A Tasmanian devil breeding program to support wild recovery. Reproduction, Fertility and Development, 31, 1296–1304.

Harrell, F.E. (2021). Hmisc: Harrell Miscellaneous. R-package v. 4.5–0.

Hauffe, H.C. & Barelli, C. (2019). Conserve the germs: the gut microbiota and adaptive potential. Conservation Genetics, 20, 19–27.

Henry, L.P., Bruijning, M., Forsberg, S.K.G. & Ayroles, J.F. (2021). The microbiome extends host evolutionary potential. Nature Communications, 12, 5141.

Hoffmann, A.A. & Nielsen, K.M. (1985). The effect of resource subdivision on genetic variation in *Drosophila*. The American Naturalist, 125, 421–430.

Hoffmann, A.A., Sgrò, C.M. & Kristensen, T.N. (2017). Revisiting adaptive potential, population size, and conservation. Trends in Ecology & Evolution, 32, 506–517.

Ingala, M.R., Becker, D.J., Bak Holm, J., Kristiansen, K. & Simmons, N.B. (2019). Habitat fragmentation is associated with dietary shifts and microbiota variability in common vampire bats. Ecology and Evolution, 9, 6508–6523.

Jaenike, J., Unckless, R., Cockburn, S.N., Boelio, L.M. & Perlman, S.J. (2010). Adaptation via symbiosis: recent spread of a *Drosophila* defensive symbiont. Science, 329, 212–215.

Jehrke, L., Stewart, F.A., Droste, A. & Beller, M. (2018). The impact of genome variation and diet on the metabolic phenotype and microbiome composition of *Drosophila melanogaster*. Scientific Reports, 8, 6215.

Kokou, F., Sasson, G., Nitzan, T., Doron-Faigenboim, A., Harpaz, S., Cnaani, A., et al. (2018). Host genetic selection for cold tolerance shapes microbiome composition and modulates its response to temperature. eLife, 7, e36398.

Koskella, B. & Bergelson, J. (2020). The study of host–microbiome (co)evolution across levels of selection. Philosophical Transactions of the Royal Society B: Biological Sciences, 375, 20190604.

Kristensen, T.N., Schönherz, A.A., Rohde, P.D., Sørensen, J.G. & Loeschcke, V. (2021). Strong experimental support for the hologenome hypothesis revealed from *Drosophila melanogaster* selection lines. bioRxiv preprint, DOI: 10.1101/2021.09.09.459587.

Leonardo, T.E. & Mondor, E.B. (2006). Symbiont modifies host life-history traits that affect gene flow. Proceedings of the Royal Society B: Biological Sciences, 273, 1079–1084.

Liang, T., Wang, X.-W., Wu, A.-K., Fan, Y., Friedman, J., Dahlin, A., et al. (2020). Deciphering functional redundancy in the human microbiome. Nature Communications, 11, 6217.

Love, M.I., Huber, W. & Anders, S. (2014). Moderated estimation of fold change and dispersion for RNA-seq data with DESeq2. Genome Biology, 15, 550.

Lozupone, C.A., Stombaugh, J.I., Gordon, J.I., Jansson, J.K. & Knight, R. (2012). Diversity, stability and resilience of the human gut microbiota. Nature, 489, 220–230.

Markert, J.A., Champlin, D.M., Gutjahr-Gobell, R., Grear, J.S., Kuhn, A., McGreevy, T.J., et al. (2010). Population genetic diversity and fitness in multiple environments. BMC Evolutionary Biology, 10, 205.

Marra, A., Hanson, M.A., Kondo, S., Erkosar, B. & Lemaitre, B. (2021). Drosophila antimicrobial peptides and lysozymes regulate gut microbiota composition and abundance, 12, 16.

McFall-Ngai, M., Hadfield, M.G., Bosch, T.C.G., Carey, H.V., Domazet-Lošo, T., Douglas, A.E., et al. (2013). Animals in a bacterial world, a new imperative for the life sciences. Proceedings of the National Academy of Sciences, 110, 3229–3236.

Memmott, J., Waser, N.M. & Price, M.V. (2004). Tolerance of pollination networks to species extinctions. Proceedings of the Royal Society of London. Series B: Biological Sciences, 271, 2605–2611.

Mistry, R., Kounatidis, I. & Ligoxygakis, P. (2017). Interaction between familial transmission and a constitutively active immune system shapes gut microbiota in *Drosophila melanogaster*. Genetics, 206, 889–904.

Moeller, A.H. & Sanders, J.G. (2020). Roles of the gut microbiota in the adaptive evolution of mammalian species. Philosophical Transactions of the Royal Society B: Biological Sciences, 375, 20190597.

Moghadam, N.N., Thorshauge, P.M., Kristensen, T.N., de Jonge, N., Bahrndorff, S., Kjeldal, H., et al. (2018). Strong responses of *Drosophila melanogaster* microbiota to developmental temperature. Fly, 12, 1–12.

Mouton, L., Henri, H., Charif, D., Boulétreau, M. & Vavre, F. (2007). Interaction between host genotype and environmental conditions affects bacterial density in *Wolbachia* symbiosis. Biology Letters, 3, 210–213.

Moya, A. & Ferrer, M. (2016). Functional Redundancy-Induced Stability of Gut Microbiota Subjected to Disturbance. Trends in Microbiology, 24, 402–413.

Muegge, B.D., Kuczynski, J., Knights, D., Clemente, J.C., Gonzalez, A., Fontana, L., et al. (2011). Diet drives convergence in gut microbiome functions across mammalian phylogeny and within humans. Science, 332, 970–974.

Mueller, U.G. & Sachs, J.L. (2015). Engineering microbiomes to improve plant and animal health. Trends in Microbiology, 23, 606–617.

Nakagawa, S., Johnson, P.C.D. & Schielzeth, H. (2017). The coefficient of determination R^2^ and intra-class correlation coefficient from generalized linear mixed-effects models revisited and expanded. Journal of The Royal Society Interface, 14, 20170213.

Oksanen, J., Blanchet, F.G., Friendly, M., Kindt, R., Legendre, P., McGlinn, D., et al. (2018). vegan: community ecology package. v 2.5-2. R-package.

Ørsted, M., Hoffmann, A.A., Sverrisdóttir, E., Nielsen, K.L. & Kristensen, T.N. (2019). Genomic variation predicts adaptive evolutionary responses better than population bottleneck history. PLOS Genetics, 15, e1008205.

Pérez-Tomé, J.M. & Toro, M.A. (1982). Competition of similar and non-similar genotypes. Nature, 299, 153–154.

Price, M.N., Dehal, P.S. & Arkin, A.P. (2009). FastTree: computing large minimum evolution trees with profiles instead of a distance Matrix. Molecular Biology and Evolution, 26, 1641–1650.

Rayes, A., Morrow, A.L., Payton, L.R., Lake, K.E., Lane, A. & Davies, S.M. (2016). A genetic modifier of the gut microbiome influences the risk of graft-versus-host disease and bacteremia after hematopoietic stem cell transplantation. Biology of Blood and Marrow Transplantation, 22, 418–422.

Redford, K.H., Segre, J.A., Salafsky, N., Rio, C.M. del & McAloose, D. (2012). Conservation and the microbiome. Conservation Biology, 26, 195–197.

Robinson, C.D., Bohannan, B.J. & Britton, R.A. (2019). Scales of persistence: transmission and the microbiome. Current Opinion in Microbiology, 50, 42–49.

Rudman, S.M., Greenblum, S., Hughes, R.C., Rajpurohit, S., Kiratli, O., Lowder, D.B., et al. (2019). Microbiome composition shapes rapid genomic adaptation of *Drosophila melanogaster*. Proceedings of the National Academy of Sciences, 116, 20025–20032.

Saleem, M., Hu, J. & Jousset, A. (2019). More than the sum of its parts: microbiome biodiversity as a driver of plant growth and soil health. Annual Review of Ecology, Evolution, and Systematics, 50, 145–168.

Sampson, T.R. & Mazmanian, S.K. (2015). Control of brain development, function, and behavior by the microbiome. Cell Host & Microbe, 17, 565–576.

Ser, H.-L., Letchumanan, V., Goh, B.-H., Wong, S.H. & Lee, L.-H. (2021). The use of fecal microbiome transplant in treating human diseases: too early for poop? Frontiers in Microbiology, 12.

Shapira, M. (2016). Gut microbiotas and host evolution: scaling up symbiosis. Trends in Ecology & Evolution, 31, 539–549.

Simhadri, R.K., Fast, E.M., Guo, R., Schultz, M.J., Vaisman, N., Ortiz, L., et al. (2017). The gut commensal microbiome of *Drosophila melanogaster* is modified by the endosymbiont *Wolbachia*. mSphere, 2, e00287-17.

Sison-Mangus, M.P., Mushegian, A.A. & Ebert, D. (2015). Water fleas require microbiota for survival, growth and reproduction. The ISME Journal, 9, 59–67.

Sommer, F. & Bäckhed, F. (2013). The gut microbiota - masters of host development and physiology. Nature Reviews Microbiology, 11, 227–238.

Spielman, D., Brook, B.W. & Frankham, R. (2004). Most species are not driven to extinction before genetic factors impact them. Proceedings of the National Academy of Sciences, 101, 15261– 15264.

Trevelline, B.K., Fontaine, S.S., Hartup, B.K. & Kohl, K.D. (2019). Conservation biology needs a microbial renaissance: a call for the consideration of host-associated microbiota in wildlife management practices. Proceedings of the Royal Society B: Biological Sciences, 286, 20182448.

Wagener, C., Mohanty, N.P. & Measey, J. (2020). The gut microbiome facilitates ecological adaptation in an invasive vertebrate. bioRxiv preprint, DOI: 10.1101/2020.12.10.418954.

Walters, A.W., Hughes, R.C., Call, T.B., Walker, C.J., Wilcox, H., Petersen, S.C., et al. (2019). The microbiota influences the *Drosophila melanogaster* life history strategy. Molecular Ecology, 29, 639–653.

Wang, S., Huang, M., You, X., Zhao, J., Chen, L., Wang, L., et al. (2018). Gut microbiota mediates the anti-obesity effect of calorie restriction in mice. Scientific Reports, 8, 13037.

Wei, L., Zeng, B., Zhang, S., Li, F., Kong, F., Ran, H., et al. (2020). Inbreeding alters the gut microbiota of the Banna minipig. Animals, 10, 2125.

Wertheim, B., Marchais, J., Vet, L.E.M. & Dicke, M. (2002). Allee effect in larval resource exploitation in *Drosophila*: an interaction among density of adults, larvae, and micro-organisms. Ecological Entomology, 27, 608–617.

West, A.G., Waite, D.W., Deines, P., Bourne, D.G., Digby, A., McKenzie, V.J., et al. (2019). The microbiome in threatened species conservation. Biological Conservation, 229, 85–98.

Wilches, D.M., Coghlin, P.C. & Floate, K.D. (2021). Next generation sequencing, insect microbiomes, and the confounding effect of *Wolbachia*: a case study using spotted-wing drosophila (*Drosophila suzukii*) (Diptera: Drosophilidae). Canadian Journal of Zoology, 99, 588–595.

Willi, Y., Kristensen, T.N., Sgrò, C.M., Weeks, A.R., Ørsted, M. & Hoffmann, A.A. (2022). Conservation genetics as a management tool: The five best-supported paradigms to assist the management of threatened species. PNAS, 119, e2105076119.

Wong, A.C.N., Vanhove, A.S. & Watnick, P.I. (2016). The interplay between intestinal bacteria and host metabolism in health and disease: lessons from *Drosophila melanogaster*. Disease Models & Mechanisms, 9, 271–281.

Yuan, M.L., Dean, S.H., Longo, A.V., Rothermel, B.B., Tuberville, T.D. & Zamudio, K.R. (2015). Kinship, inbreeding and fine scale spatial structure influence gut microbiota in a hindgut fermenting tortoise. Molecular Ecology, 24, 2521–2536.

Zilber-Rosenberg, I. & Rosenberg, E. (2008). Role of microorganisms in the evolution of animals and plants: the hologenome theory of evolution. FEMS Microbiology Reviews, 32, 723–735.

